# MMpred: a distance-assisted multimodal conformation sampling for de novo protein structure prediction

**DOI:** 10.1101/2021.01.21.427573

**Authors:** Kai-Long Zhao, Jun Liu, Xiao-Gen Zhou, Jian-Zhong Su, Yang Zhang, Gui-Jun Zhang

**Affiliations:** College of Information Engineering, Zhejiang University of Technology, Hangzhou 310023, China; Department of Computational Medicine and Bioinformatics, University of Michigan, 100 Washtenaw, Ann Arbor, MI 48109-2218, USA; School of Biomedical Engineering, School of Ophthalmology and Optometry and Eye Hospital, Wenzhou Medical University, Wenzhou 325011, Zhejiang, China

## Abstract

**Motivation:** The mathematically optimal solution in computational protein folding simulations does not always correspond to the native structure, due to the imperfection of the energy force fields. There is therefore a need to search for more diverse suboptimal solutions in order to identify the states close to the native. We propose a novel multimodal optimization protocol to improve the conformation sampling efficiency and modeling accuracy of de novo protein structure folding simulations.

**Results:** A distance-assisted multimodal optimization sampling algorithm, MMpred, is proposed for de novo protein structure prediction. The protocol consists of three stages: The first is a modal exploration stage, in which a structural similarity evaluation model DMscore is designed to control the diversity of conformations, generating a population of diverse structures in different low-energy basins. The second is a modal maintaining stage, where an adaptive clustering algorithm MNDcluster is proposed to divide the populations and merge the modal by adjusting the annealing temperature to locate the promising basins. In the last stage of modal exploitation, a greedy search strategy is used to accelerate the convergence of the modal. Distance constraint information is used to construct the conformation scoring model to guide sampling. MMpred is tested on a large set of 320 non-redundant proteins, where MMpred obtains models with TM-score≥0.5 on 268 cases, which is 20.3% higher than that of Rosetta guided with the same set of distance constraints. The results showed that MMpred can help significantly improve the model accuracy of protein assembly simulations through the sampling of multiple promising energy basins with enhanced structural diversity.

**Availability:** The source code and executable versions are freely available at https://github.com/iobio-zjut/MMpred.

## 1 Introduction

As the function of a protein is dependent on its structure, predicting protein structure from its amino acid sequence has been a grand challenge in biology for decades (Abriata *et al*., 2019; Bradley *et al*., 2005; Dill and MacCallum, 2012). Protein structure prediction, especially de novo prediction, has made a major breakthrough in recent years, due to the drastic improvement of the accuracy of residue-residue contact and distance prediction (Shrestha *et al*., 2019). Nevertheless, due to the high dimensionality of protein conformational space, conformation sampling represents still one of the main bottlenecks in de novo protein structure prediction. Monte Carlo (Zhang et al., 2002; Rohl *et al*., 2004; Gregory *et al*., 2009; Xu and Zhang 2012), evolutionary algorithm (Zhou *et al*., 2019; Liu *et al*., 2019; Peng *et al*., 2020), multi-objective optimization (Olson and Shehu, 2014), and many other sampling strategies have been proposed to explore the conformational space.

Large-scale conformational sampling is usually based on coarse-grained energy function (Kuhlman *et al*., 2019; Francois *et al*., 2011). However, the coarse-grained model also introduces inaccuracy while reducing the search space dimension and improving computational efficiency (Lindorff-Larsen *et al*., 2011; Lazaridis and Karplus, 2000). As illustrated in **Figure 1**, the lowest energy model (in basin A) does not necessarily correspond to the native structure, and the model in the local basin B of the energy landscape may be closer to it. Therefore, searching only the lowest free energy structure is insufficient, and sampling local minimum solutions with structural diversity in different low energy basins is also important (Park *et al*., 2018; Rohl *et al*., 2004; Zhou *et al*.,2019). Rosetta samples a large number of low-energy conformations by running thousands of independent Metropolitan Monte Carlo trajectories in parallel to reveal the local minimum of the energy surface (Raman *et al*.,2008; Zhou and Zhang, 2018). C-QUARK performs a replica-exchange Monte Carlo folding simulation to generate a large number of conformations, which are submitted to SPICKER for clustering, and the model is selected in the most populated clusters instead of the clusters with lowest energy (Zheng *et al*., 2019; Xu and Zhang, 2012). Memetic evolutionary algorithm has been proposed to sample a diverse ensemble of conformations through genetic operators, which promote the exploration of protein conformational space (Garza-Fabre *et al*., 2016). These methods alleviate the sampling problem based on inaccurate energy function to a certain extent, but the adaptive determination of cluster number and rapid evaluation of structural similarity need further to be investigated.

**Fig. 1.**
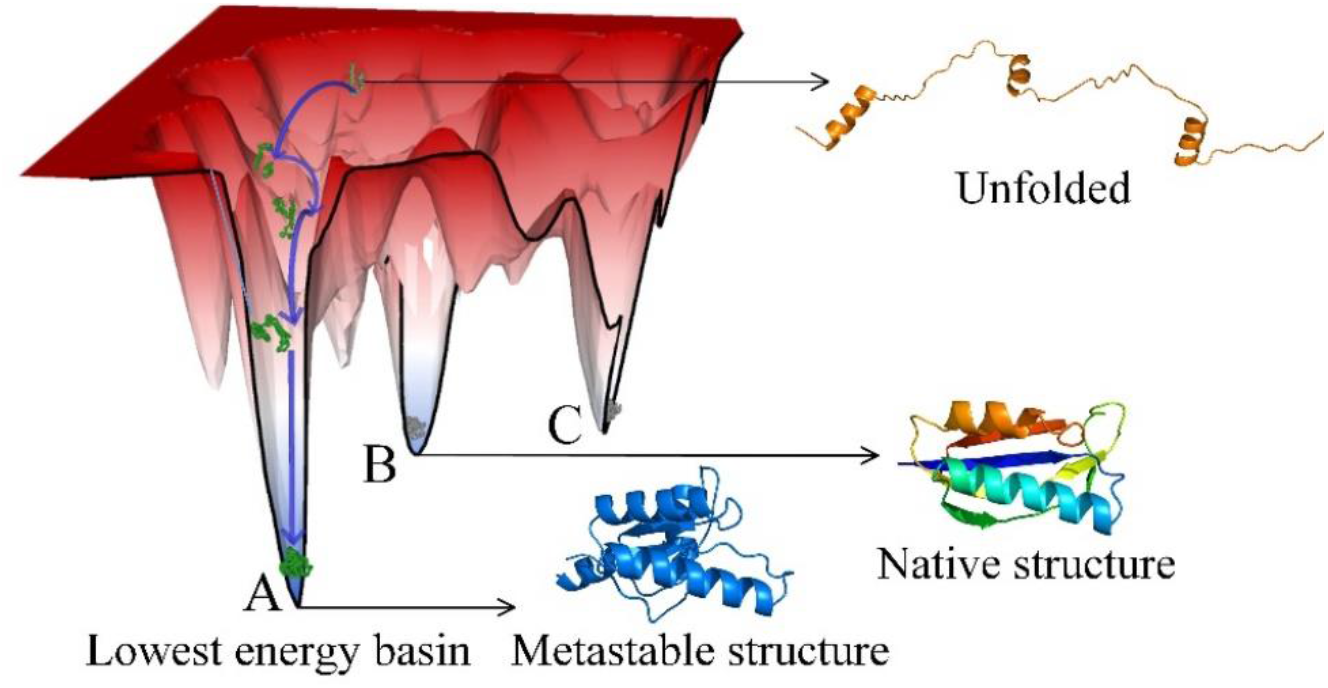
Schematic diagram of protein-folding energy landscape.

To help reshape the energy landscape, inter-residue contact and distance maps have been proposed and shown usefulness for de novo protein structure prediction (Senior *et al*., 2019; Xu and Wang, 2019; Kandathil *et al*., 2019; Li *et al*., 2019). In particular, AlphaFold and RaptorX focused on distance prediction and distancebased protein structure modeling in CASP13. AlphaFold (A7D) constructs the potential energy of the average force from the predicted distance information, which is optimized by the stochastic gradient descent algorithm to generate the structure (Senior *et al*., 2020). RaptorX inputs the predicted distance, secondary structure, and torsion angles predicted by deep convolutional neural networks into the Crystallography and NMR system (CNS) (Brunger et al., 2007) to quickly constructed an accurate 3D model (Xu, 2019; Wang *et al*., 2017). In DMPfold (Gregory *et al*.,2019), the inter-residue distances, hydrogen bonds and torsion angles predicted from DMP are used to generate models with CNS, and a single model is used as additional input to refine the distances and hydrogen bonds. trRosetta introduces inter-residue orientation prediction and uses the predicted orientation and distance to construct constraints for Rosetta-constrained energy minimization protocol in order to generate structure models (Yang *et al*., 2020). However, the predicted contact distance is not sufficiently accurate and has the same defects as the abovementioned energy function (Hou *et al*., 2019).

De novo protein structure prediction based on energy (or distance) -guided conformation sampling is essentially an uncertain optimization problem. Multimodal optimization can discover and maintain multiple feasible solutions (Stoean *et al*., 2010; Sareni *et al*., 1998), which is critical for improving the efficiency of conformational sampling. In this study, a distance-assisted de novo protein structure prediction by multimodal conformation sampling algorithm (MMpred) is proposed. First, the crowding strategy is designed based on the structural similarity evaluation model to explore the conformation space extensively and generate a population of conformations with structural diversity. Then, an adaptive clustering algorithm and a modal merging strategy are used to divide and maintain the modal, respectively. Finally, a greedy search strategy is used to accelerate the convergence of the modal. MMpred can discover and exploit multiple promising low-energy basins to sample a set of diverse structures and ultimately improve structure prediction accuracy.

## 2 Materials and methods

The pipeline of MMpred is illustrated **Figure 2**. For the query sequence, the fragment library is built by Robetta (http://robetta.bakerlab.org/), and the inter-residue distance is predicted by trRosetta (https://yanglab.nankai.edu.cn/). The initial population is generated by the random fragment assembly. The population goes through three stages of evolution, and finally five prediction models are generated. In the first modal exploration stage, a structural similarity evaluation model, DMscore, is designed to select the conformation that is the most similar to the trial conformation produced by fragment assembly in the population for conformation replacement. In the second modal maintaining stage, an adaptive clustering algorithm, MNDcluster, is proposed to divide the population and then merge the modal by adjusting the annealing temperature to locate promising models. In the final modal exploitation stage, a greedy search strategy is used to accelerate the convergence of the modal. Finally, the modal are sorted according to the average distance score of the conformations in the modal, and the model with the lowest distance score among the top five modals is selected as the final model.

**Fig. 2.**
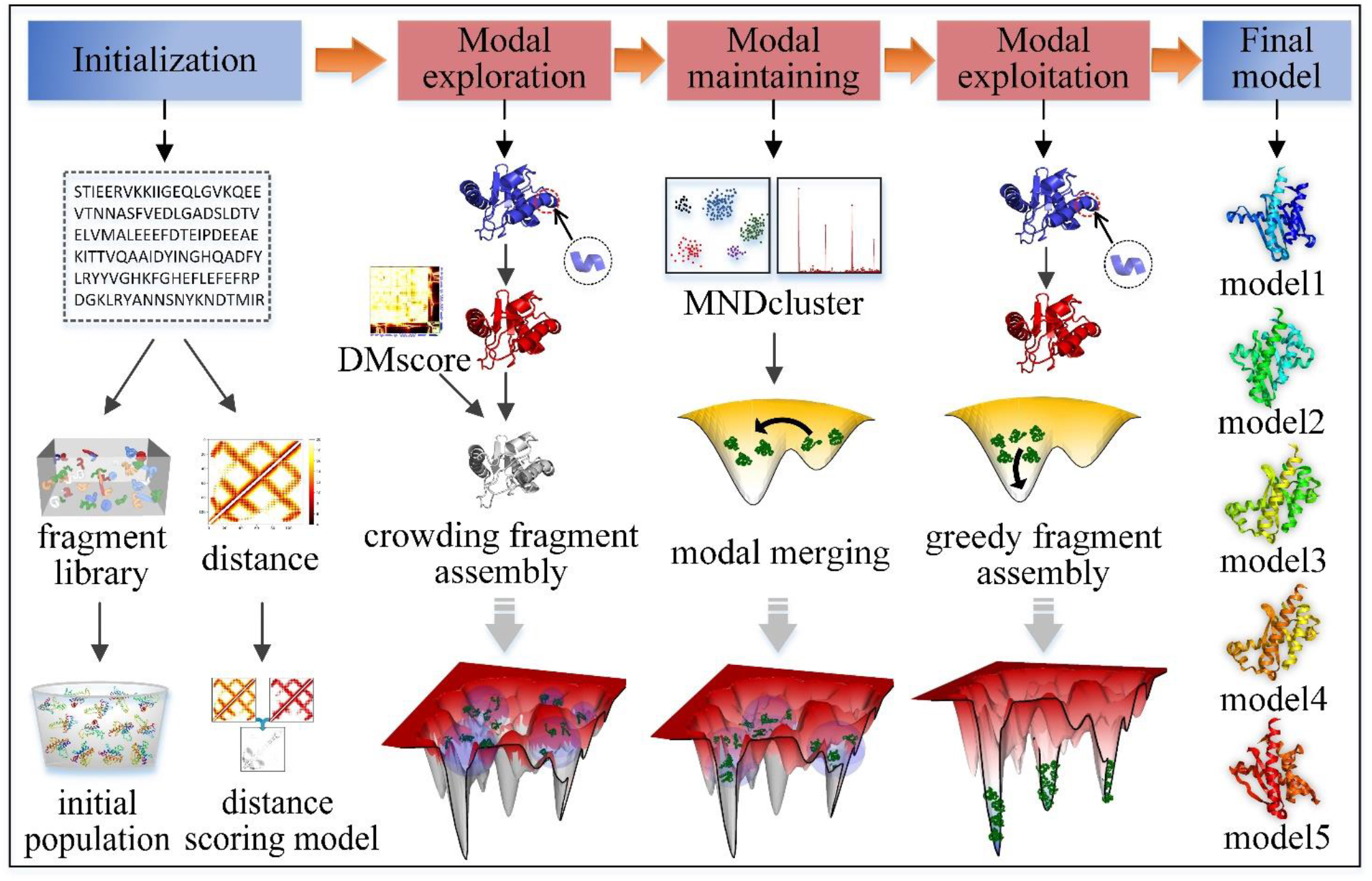
The pipeline of MMpred for multimodal conformation sampling.

### 2.1 Modal exploration

Maintaining the diversity of the population during the evolution is the key to the multimodal optimization algorithm. The purpose of modal exploration is to generate a conformation population with structural diversity in different low-energy regions through extensive sampling of conformational spaces to obtain a broad view. We use a crowding strategy to maintain population diversity during evolution (De Jong, 1975; Ling *et al*., 2008). Its principle is to replace the most similar conformation in the population with an offspring. However, effective crowding strategies depend on similarity metrics (Thomsen *et al*., 2004). Therefore, we designed a structural similarity evaluation model based on the comparison of residue distance maps (DMscore) to evaluate the structural similarity between two conformations (Simoncini *et al*., 2017; Zhang *et al*., 2004; Cheng *et al*., 2019).

#### 2.1.1 DMscore

DMscore measures the similarity by calculating the difference of Euclidean distance of the corresponding residue pairs in two models, which is defined as follows:

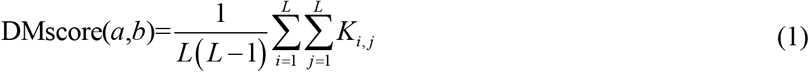

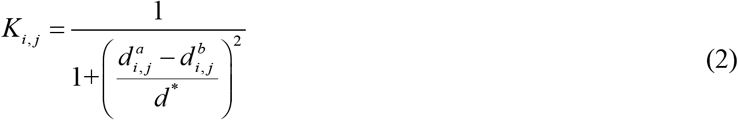

where *L* is the length of the protein structure; 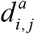 and 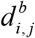. are the distances between the *i* th and *j* th residues in model *a* and model *b*, respectively; and *d*\* is the normalized scale used to eliminate the inherent dependence of the score on protein size. The value range of 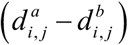 varies with the sequence distance between *i* and *j*. So it is unreasonable to set *d*\* as a constant. We analyzed the relationship between the average Euclidean distance between residues and the length of the protein on the native structure of 500 non-redundant targets with length ranges from 40 to 700. As shown in **Supplementary Figure S1**, the average distance between residues increases logarithmically with increasing protein length. Therefore, *d*\* is defined as the logarithm of the residue sequence distance, namely:

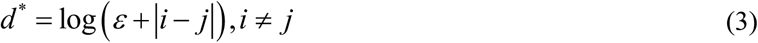

where *ε* is an infinitely small quantity to avoid *d*\* being zero. The value of DMscore is between (0,1]. The higher value indicates higher similarity between the two models; 1 indicates a perfect match between two models.

Without complex structure rotation and translation such as RMSD and TM-score (Zhang et al., 2004), DMscore can quickly evaluate the structural similarity of two conformations by calculating the difference of Euclidian distance of corresponding residual pairs. As shown in **Figure 3**, the comparison of distance maps can well characterize the similarity of two structures. The structural similarity measure which is based on the global rigid body superposition has limitations. The similarity score is dominated by the part with the highest coverage. So, the part with the lowest coverage cannot be matched correctly, leading to artificially unfavorable scores (Hubbard *et al*., 1999; Mariani *et al*., 2013). DMscore not only evaluates the global topology but also considers the local structure differences.

**Fig. 3.**
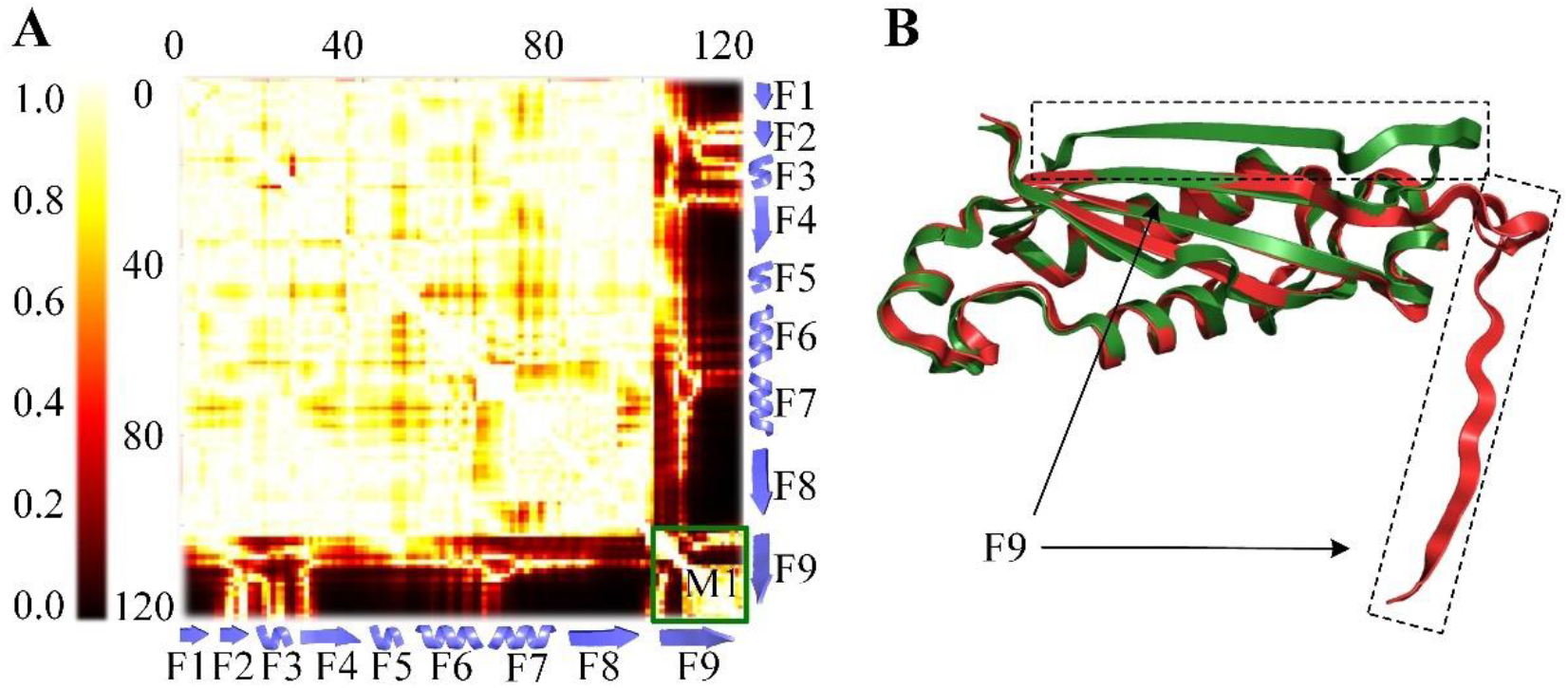
(A) is the comparison of the real distance map of two models, each point represents *K_i,j_*; (B) is the superposition of the two models. The F1 to F8 of the two models in (B) are well aligned, corresponding to the bright color area (indicating high scores) in the upper left part of (A). F9 deviates from the overall structure significantly. The distance between F9 and other fragments in the two models is significantly different and corresponds to the dark area, which indicates low score. However, the F9 fragments of the two models are very similar. M1 shows a light color in the map, which gives a high score to local fragments.

#### 2.1.2 Distance scoring model

The predicted distance constraint is used to construct the distance scoring model, which is used to guide conformational sampling along with the energy function (Rosetta score3) (Ovchinnikov *et al*., 2018). The principle of the conformation scoring model based on the predicted inter-residue distance, named *D_score_*, is the same as DMscore, as shown in **Supplementary Figure S2**. The difference is that *D_score_* scores the conformation by calculating the similarity between the real distance map of the conformation and the predicted distance map. *D_score_* can be calculated by

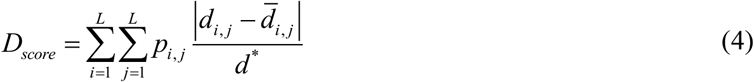

where *d_i,j_* and 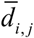 are the distances between the *i*th and *j*th residues in target conformation and predicted distance map, respectively; *d*\* is a normalized scale calculated by Equation (3). *P_i,j_* is the probability of predicted distance. The distance difference is not calculated for the residue pair with no predicted value of distance. A smaller *D_score_* indicates the conformation satisfies better the predicted distance constraints.

#### 2.1.3 Crowding fragment assembly

The crowding fragment assembly is shown in **Supplementary Figure S4**. The trial conformation is obtained by the fragment assembly of the target conformation. The similarity between the trial conformation and each one in the population was calculated by DMscore. In order to improve the success rate of crowding, the top *R* conformations with the highest DMscore to the trial conformation are selected as the crowded subpopulation. The trial conformation is then attempted sequentially to replace each of the top *R* conformations starting with the ones with the highest DMscore until the replacement is successful or traverses all conformations in the subpopulation. Conformation replacement is performed by the Metropolis criterion based on energy function and *D_score_* sequentially. After successive iterations, a population with diverse structures in different low-energy region is generated.

### 2.2 Modal maintaining

After obtaining a population with diverse structure distribution, the population will be gradually divided into multiple promising modals in the maintaining stage. Considering the differences in the population distribution of different target proteins, we propose a clustering algorithm to adaptively determine the number of clusters according to the multi-order nearest distance analysis (MNDcluster) (Rodriguez *et al*., 2014). MNDcluster can avoid unreasonable population division caused by a fixed number of clusters. Modal exploration may result in the conformation falling into the unpromising basins. So, we designed a modal merge strategy to merge the unpromising clusters and maintain the promising clusters by adjusting the annealing temperature.

#### 2.2.1 MNDcluster

MNDcluster can obtain the global distribution information of the population by analyzing the distance of multi-order nearest neighbours. The number of clusters is adaptively counted by step information. Then, the K-medoids (Kaufman and Rousseeuw, 2009) algorithm is used to cluster the conformations according to the number of clusters.

The DMscore between conformations in the population is taken as their distance. 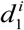 is the first-nearest neighbour distance of the *i* th conformation in the population, 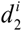 is the second-nearest neighbour distance, and so on. These distances satisfy:

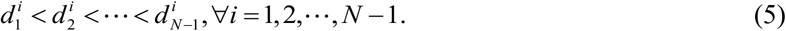

where *N* is the population size. The mean value and the mean square value of the *j* th nearest neighbour distances of all conformations are calculated as follows:

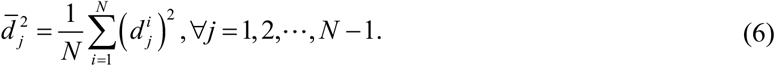

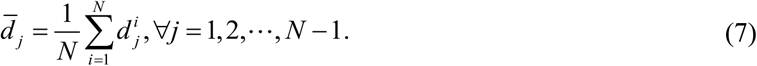

The distribution of individuals determines the number of divisible populations. The distribution degree of the population is analyzed by calculating the variance, as follows:

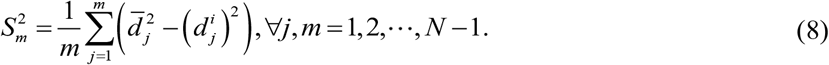

As shown in **Supplementary Figure S3**, the curvature change of variance linearized among individuals in the population is analyzed. The slope of the neighbouring individuals of the same subpopulation varies little. When the slope is calculated from one subpopulation to another, it fluctuates greatly. In the case of multiple populations, all step-jumping information can be captured in this way, and the global distribution of the current population can be obtained. MNDcluster can determine the distribution of the whole population without prior information and then obtain the statistics of the divisible population.

#### 2.2.2 Modal merging

In order to identify the promising modal, a modal evaluation model based on cluster density and cluster size is designed as:

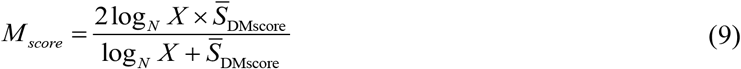

where *N* is the population size; *X* is the cluster size; and 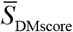 is the average DMscore between conformations in the cluster that reflects the density of the cluster. We define clusters with *M_score_* greater than or equal to *λ* as promising modals.

In the modal maintaining stage, Monte Carlo simulated annealing search and crowding strategy are used to update the modal. The annealing temperature is adjusted to maintain the promising modalities and merge the unpromising modalities. The Monte Carlo simulated annealing search can accept the conformation with increased energy according to the probability. Its dynamic properties enable the conformation to jump out of the energy trap. The conformation update probability can be calculated by:

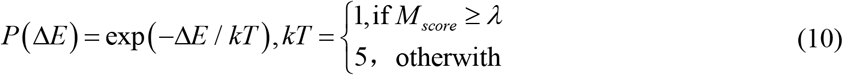

where Δ*E* is the energy difference between the target conformation and trial conformation; and *kT* is the artificial “temperature” (Li and Scheraga, 1987). For promising modal, *kT* is set to 1. The purpose is to maintain the promising modal and prevent the loss of the modal caused by the deviation of the conformation from the current basin. For the unpromising modal, *kT* is set to 5. High temperature can increase the probability of conformation update. Thus, the conformation jumps out of the current basin and enters a promising basin.

The conformational update during the modal maintaining stage is shown in **Supplementary Figure S5**. The DMscore between the trial conformation of the unpromising modal and the centroid of the nearest promising modal is calculated. The two conformations are considered to have similar topologies when DMscore is greater than or equal to 0.5, and the trial conformation is merged into the promising modal.

### 2.3 Modal exploitation

In the modal exploitation stage, the explored promising modal rapidly converge to the minimum by greedy searching, the flowchart is shown in **Supplementary Figure S6**. The target conformation is randomly selected from the subpopulation. The trial conformation is obtained by fragment assembly of the target. The distance scoring model and energy scoring model are successively used to evaluate the trial conformation. Metropolis criterion is used to determine whether the conformation with the lowest energy or the lowest *D_score_* is replaced by trial conformation in the subpopulation. If it failed, the target conformation is passed on to the next generation. The greedy search strategy converges the subpopulations to the minimum of multiple basins, and then, a diverse set of protein conformations is sampled.

## 3 Results

### 3.1 Datasets

The benchmark set is constructed according to the SCOPe 2.07 (Fox *et al*., 2014). CD-HIT (Fu *et al*., 2012) is used to cluster the SCOPe dataset with a sequence identity cut-off of 30%, and the representative proteins of each cluster are selected to form 11,198 proteins. The protein is discarded if its length is outside the range of 50-200 residues or contains multiple domains, which result in 2,481 proteins. Finally, 320 proteins are randomly selected as the benchmark set by considering the protein type and length diversity. The detailed information of the benchmark set is shown in **Supplementary Table S1**. In addition, the 24 free modeling (FM) targets of the CASP13 experiment are used to further test the performance of MMpred. The detailed information is shown in Supplementary **Table S2**. The parameters of MMpred are described in **Table 1**.

**Table 1.** The parameter descriptions in MMpred.

|  |  |  |
| --- | --- | --- |
| Population size | $N$ | 400 |
| Generation of exploration stage | $G_a$ | 500 |
| Generation of maintaining stage | $G_b$ | 300 |
| Generation of exploitation stage | $G_c$ | 800 |
| Crowded subpopulation size (exploration) | $R_a$ | $N/20$ |
| Crowded subpopulation size (maintaining) | $R_b$ | $N/40$ |
| Energy temperature scaling factor | $KT$ | 2 |
| $D_{score}$ temperature scaling factor | $KT_d$ | 2 |
| $M_{score}$ threshold | $\lambda$ | 0.5 |
| infinitely small quantity | $\varepsilon$ | 0.001 |

### 3.2 Comparison of MMpred and Rosetta-distance

MMpred is compared with distance-constrained Rosetta (Rosetta-distance) on the benchmark set. Both use the same fragment library without homologous structure. For Rosetta-distance, the distance scoring model that is the same as MMPred is used to guide the fragment assembly together with the energy function. 400 independent trajectories are run using Rosetta’s ClassicAbinitio protocol with the increase_cycles equal to 1. The central model of the first cluster determined by SPICKER (Zhang and Skolnick, 2004) is considered as the final model.

The predicted results of MMpred and Rosetta-distance on the benchmark set are summarized in **Table 2**, and the detailed results of each protein are presented in **Supplementary Table S1**. The average TM-score of the first model by MMpred (0.629) is 17% higher than that of Rosetta-distance (0.537), and the average RMSD of MMpred (6.63Å) are reduced by 12.8% compared to Rosetta-distance (7.48Å). MMpred and Rosetta- distance obtain models with TM-score≥0.5 on 268 and 203 out of 320 proteins, accounting for 84% and 63% of the total protein, respectively. The significance test results (*p*-value = 1.09E-45) show that the performance of MMpred is significantly better than that of Rosetta-distance. Since MMpred and Rosetta-distance use the same distance constraint and fragment library, the comparison results reflect the contribution of multimodal optimization to the improvement of prediction accuracy. **Figure 4A** shows the TM-score distribution of the first model predicted by MMpred and Rosetta-distance.

**Fig. 4.**
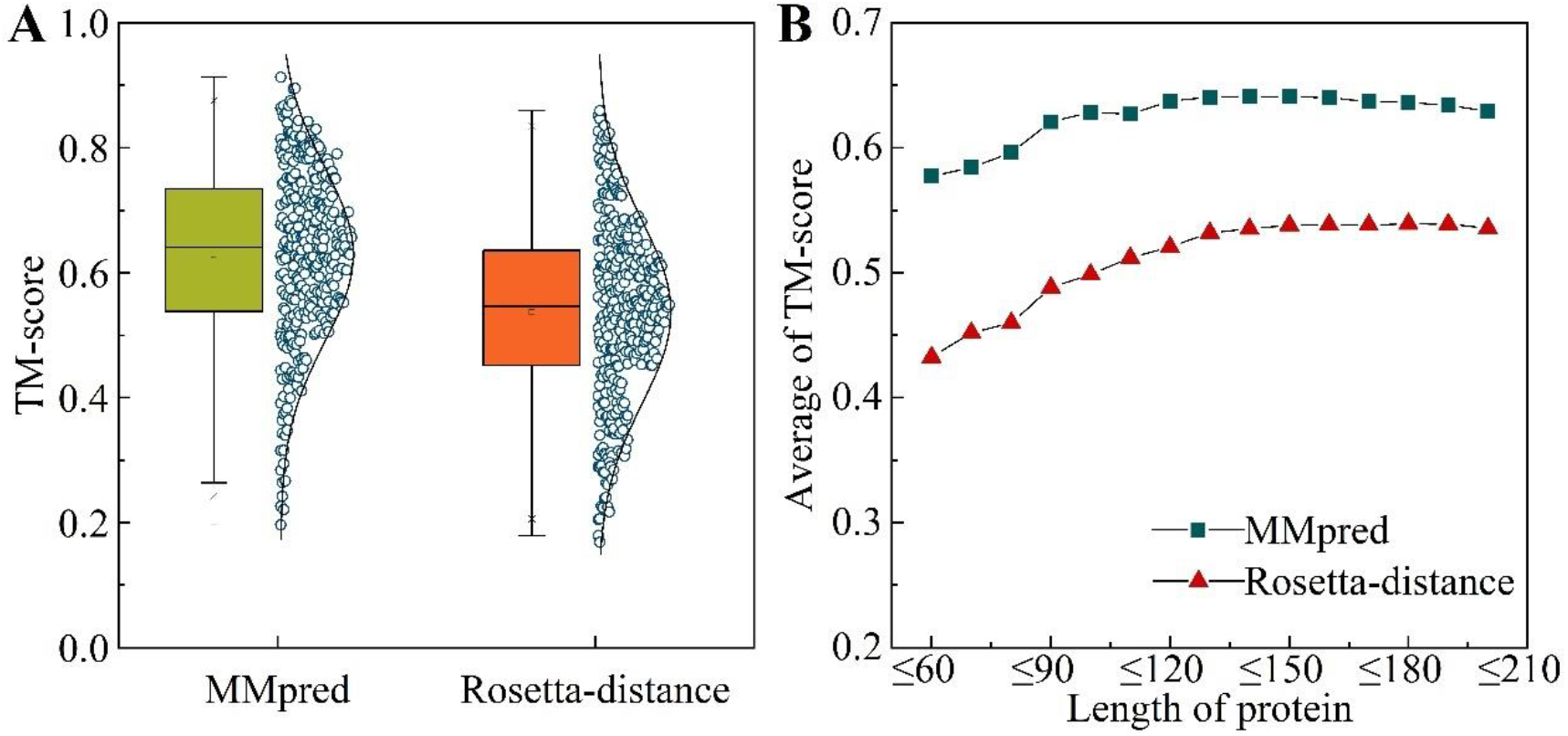
(A) Mean and distribution of TM-score for first model by MMpred and Rosetta-distance. (B) The relationship between the average TM-score of the first model of the benchmark set and the sequence length. The x-axis represents the threshold of sequence length, and the y-axis represents the average TM-score of all predicted models whose sequence length is less than the threshold.

**Table 2.**
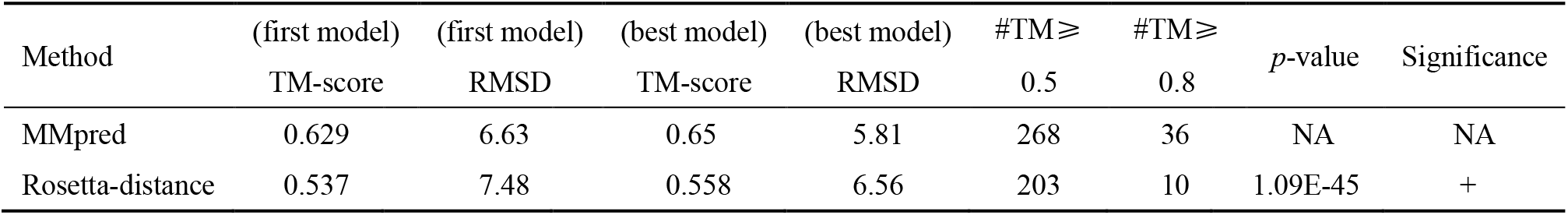
The predicted results of MMpred and Rosetta-distance. #TM ≥ 0.5 and #TM ≥ 0.8 are the number of the first model with TM-score ≥ 0.5 and TM-score ≥ 0.8, respectively. The last two columns are the results of the Wilcoxon signed rank test calculated based on the TM-score of the first model.

| Method | (first model)<br>TM-score | (first model)<br>RMSD | (best model)<br>TM-score | (best model)<br>RMSD | #TM≥<br>0.5 | #TM≥<br>0.8 | $p$-value | Significance |
| --- | --- | --- | --- | --- | --- | --- | --- | --- |
| MMpred | 0.629 | 6.63 | 0.65 | 5.81 | 268 | 36 | NA | NA |
| Rosetta-distance | 0.537 | 7.48 | 0.558 | 6.56 | 203 | 10 | 1.09E-45 | + |

**Figure 4B** shows the relationship between the TM-score and the sequence length. When the threshold of sequence length is less than or equal to 150, the average TM-score of MMpred gradually increases. The maximum value of 0.641 is reached when the threshold is equal to 150. With the further increase of the threshold value, the average TM-score decreases slightly. Part of the reason for the decreased performance is that the protein conformation space expands exponentially with the increase of sequence length. Therefore, there are more deceptive traps for proteins with larger sequence length, and MMpred needs to generate more conformational samples for the larger-size proteins to explore more promising low-energy basins. **Supplementary Figure S7** shows an example of a protein (PDB ID: 1RZ3_A) with the sequence length of 184. The TM-score of the predicted model increases from 0.463 to 0.607 when the population size increases from 400 to 800 and the generation of modal exploitation increases from 800 to 2000.

### 3.3 Results of CASP13 test set

MMpred is compared with four CASP server groups, i.e., C-QUARK (Zheng *et al*., 2019), RaptorX-DeepModeller (Xu and Wang, 2019), BAKER-ROSETTASERVER (Park *et al*., 2019) and MULTICOM_CLUSTER (Hou *et al*., 2019) on 24 FM targets of CASP13. The results of these four groups are from the CASP official website (https://predictioncenter.org/download_area/). The detailed results of MMpred and these four methods are shown in **Supplementary Table S2**. MMpred obtains the highest TM-score on 7 targets, while C-QUARK, RaptorX-DeepModeller, BAKER-ROSETTASERVER, and MULTICOM_CLUSTER get the highest TM-score on 5, 8, 3, and 1 protein(s), respectively. MMpred achieves an average TM-score of 0.466 on all targets, which is slightly lower than those of C-QUARK and RaptorX-DeepModeller but higher than those of BAKER-ROSETTASERVER and MULTICOM_CLUSTER. The accuracies of the MMpred models on the three targets (T1005-D1, T0950-D1 and T0969-D1) with a sequence length greater than 300 are relatively low. **Figure 5** shows the relationship between model accuracy and protein size. When the sequence length is less than 300, little difference exists between MMpred and C-QUARK, and RaptorX-Deepmodeller. When the sequence length is less than 150 (with 16 proteins), MMpred achieves an average TM-score of 0.493, which is slightly higher than that by C-QUARK (0.488) and RaptorX-Deepmodeller (0.463).

**Fig. 5.**
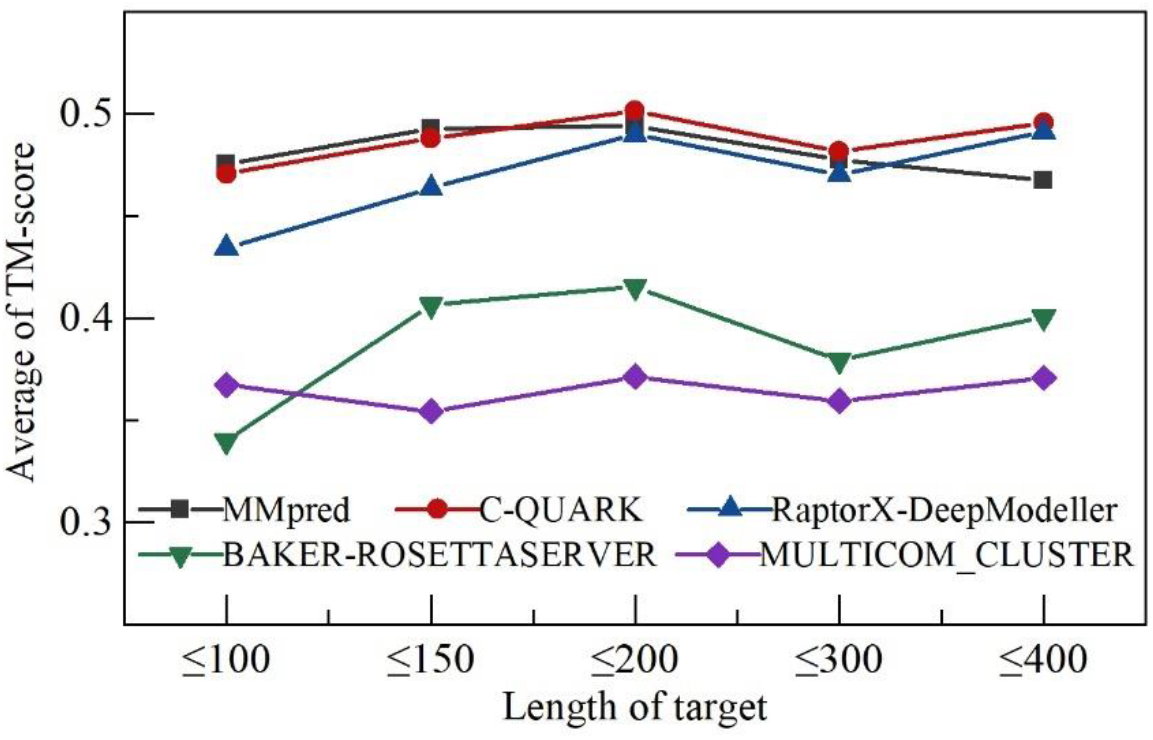
The relationship between the average TM-score of the first model and the sequence length on the 24 FM targets of CASP13. The x-axis represents the threshold of sequence length, and the y-axis represents the average TM-score of all predicted models whose sequence length is less than the threshold.

To further test the prediction performance of MMpred, a comparison with C-QUARK was performed on the 320 benchmark proteins. The result of C-QUARK is predicted by its online server (https://zhanglab.ccmb.med.umich.edu/C-QUARK/). C-QUARK uses multiple contact-maps and performs replica-exchange Monte Carlo for protein folding. The predicted results of MMpred and C-QUARK on the benchmark set are summarized in **Table 3**, and the detailed results of each protein are presented in **Supplementary Table S1**. The average TM-scores of the first model by MMpred and C-QUARK are 0.629 and 0.623, respectively. MMpred and C-QUARK generate models with TM-score≥0.5 on 268 and 256 proteins, accounting for 84% and 80% of the total protein, respectively. The significance test results (*p*-value = 0.723) show that the performance of MMpred is comparable with that of C-QUARK.

**Table 3.** The predicted results of the first model of MMpred and C-QUARK. #TM ≥ 0.5 is the number of the first model with TM-score ≥ 0.5. The last two columns are the results of the Wilcoxon signed rank test calculated based on the TM-score of the first model.

| Method | TM-score | #TM≥0.5 | $p$-value | Significance |
| --- | --- | --- | --- | --- |
| MMpred | 0.629 | 268 | NA | NA |
| C-QUARK | 0.623 | 256 | 0.723 | ≈ |

### 3.4 Component analysis

To examine the effect of the three stages of MMpred on the performance of the algorithm, we construct and compare three different versions of MMpred, MMpred-A_C, MMpred-B_C and MMpred-C. MMpred- A_C includes the modal exploration and the modal exploitation, MMpred-B_C uses the modal maintaining and the modal exploitation, and MMpred-C only contains the modal exploitation. The predicted results of MMpred, MMpred-A_C, MMpred-B_C and MMpred-C on the benchmark set are summarized in **Table 4**, and the detailed results of each protein are presented in **Supplementary Table S3**.

**Table 4.** The predicted results of the first model of MMpred, MMpred-A_C, MMpred-B_C and MMpred-C. #TM ≥ 0.5 is the number of the first model with TM-score ≥ 0.5. The last two columns are the results of the Wilcoxon signed rank test calculated based on the TM-score of the first model.

| Method | TM-score | #TM ≥ 0.5 | *p*-value | Significance |
| --- | --- | --- | --- | --- |
| MMpred | 0.629 | 268 | NA | NA |
| MMpred-A_C | 0.596 | 197 | 1.13E-11 | + |
| MMpred-B_C | 0.541 | 241 | 3.94E-41 | + |
| MMpred-C | 0.534 | 189 | 3.13E-42 | + |

Compared with MMpred-C, the average TM-score of the first model and the best model of MMpred-A_C improved by 11.5% and 11.6%, respectively. This shows that modal exploration is very important. It can discover and develop more potential basins and generate populations with higher structural diversity. The prediction accuracy of MMpred-B_C is only slightly improved compared with MMpred-C. This is due to the lack of effective exploration in the early stage, resulting in the modal maintaining stage cannot play an effective role. This can be reflected in the comparison with MMpred. After adding the exploration stage, the average prediction accuracy increases from 0.541 to 0.629, with an increase of 16.3%. The number of models with TM-score≥0.5 also increases by 71. Compared with MMpred-A_C, the prediction accuracy of MMpred with modal maintaining stage added has also significantly improved. The average prediction accuracy is improved by 5.5%, and the number of models with TM-score≥0.5 is increased by 11.2%. The results of the significance test show that the prediction accuracy of MMpred is significantly better than each of its comparative versions. **Figure 6** visually reflects the comparison between MMpred and each comparison method in the first model and in the best model. MMpred outperforms the individual programs (MMpred-A_C, MMpred-B_C and MMpred-C) in 198, 264 and 274 cases when considering the first models. This data demonstrate again that the importance of the different step of multimodal optimization.

**Fig. 6.**
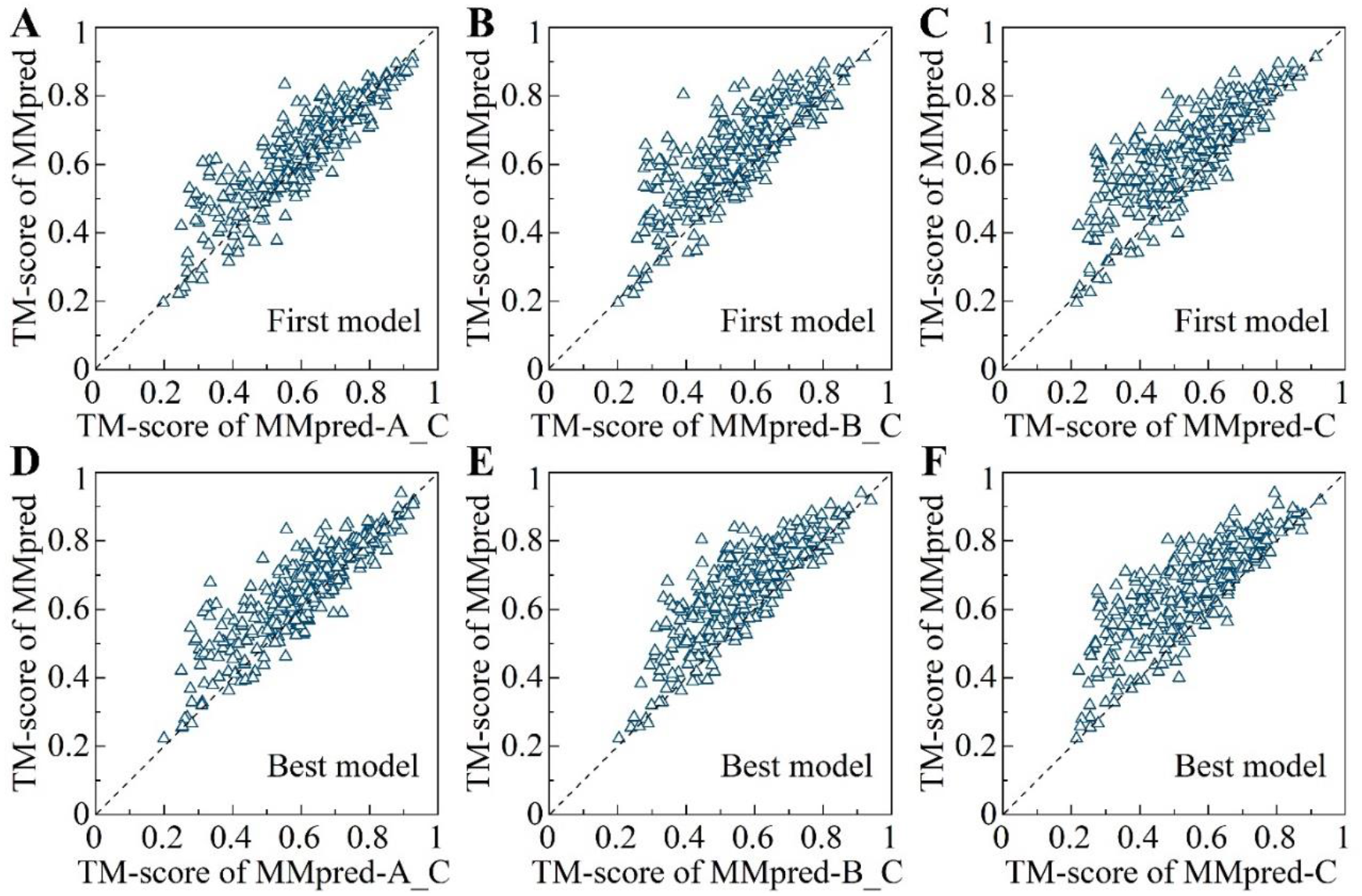
The comparison of MMpred with MMpred-A_C, MMpred-B_C, and MMpred-C. (A-C) are the TM-score of the predicted first model between MMpred and MMpred-A_C, MMpred-B_C, and MMpred-C, respectively. (D-F) are the TM-score of the predicted best model of between MMpred and MMpred-A_C, MMpred-B_C, and MMpred-C, respectively.

In order to verify the role of the crowding strategy in the modal exploration stage, 40 proteins were randomly selected from the benchmark set for analysis. We respectively calculated the diversity of the population obtained by using the crowding strategy and by the method without crowding strategy in the modal exploration stage. The results are shown in **Figure 7**. The diversity of the population is characterized by the average value of the DMscore of any two individuals in the population. A lower DMscore indicates a greater population diversity. The results indicate that the use of crowding strategy in all proteins can better maintain the diversity of the population.

**Fig. 7.**
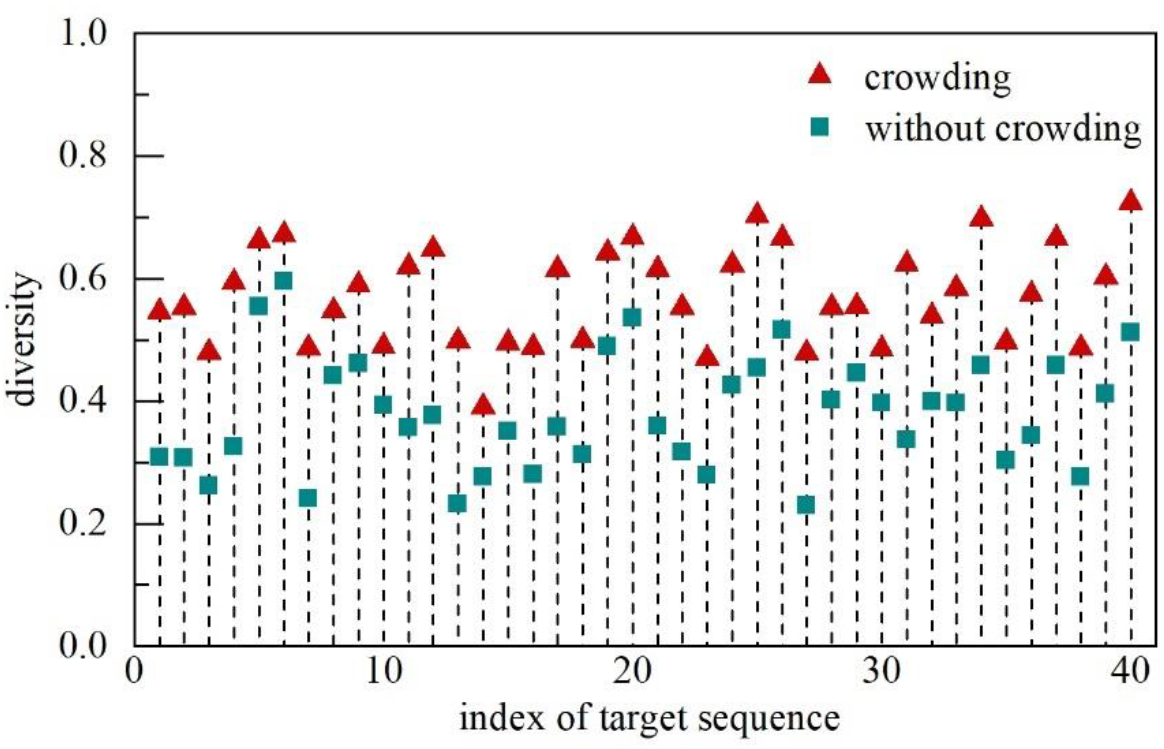
Comparison of population diversity between the method using crowding strategy and that without crowding strategy in the modal exploration stage.

### 3.5 Experimental analysis of DMscore

DMscore is tested on 20 targets of CASP13, each containing the first model predicted by all the server groups. For each target, we calculated the DMscore between the first model predicted by all groups and its native structure and analyzed its correlation with GDT_TS. Then, we compared it with TM-score. The comparison results of all proteins are shown in **Supplementary Figure S8**. The DMscore and GDT_TS have a strong correlation on the predicted models by different groups, especially for the model with DMscore≥0.5. The characteristic of DMscore is that it can quickly evaluate the similarity between models without rotation and translation, and it focuses on topological structure matching. **Figure 8** shows an example of T0954-D1. For the 78th model ranked by GDT_TS, the TM-score is 0.31, but DMscore is 0.74. By observing the 75th to 78th models and the native structure, the topological structure of the 75th to 77th models is disordered, and the topological structure of the 78th model is more similar to the native structure and may be better.

**Fig. 8.**
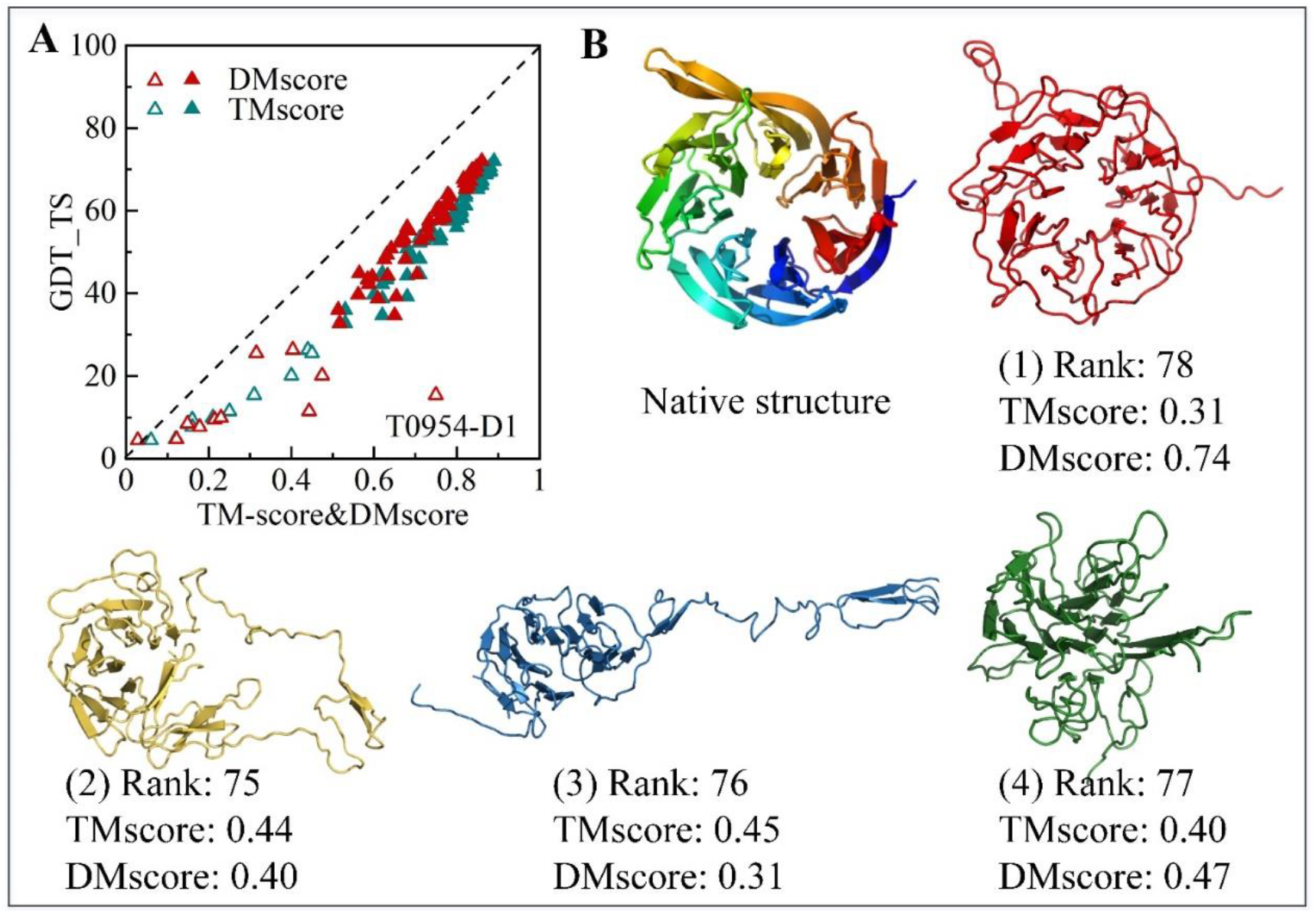
Example of DMscore analysis of CASP13 target T0954-D1. (A) Comparison of the relationship among DMscore, TM-score and CDT_TS of the first model predicted by all groups. The model with DMscore or TM-score greater than or equal to 0.5 is represented as a solid triangle, and the 78th model ranked by GDT_TS is marked. (B) The native structure and the structure of the 75th to 78th models.

### 3.6 Case studies

**Figure 9** shows the RMSD-energy scatter diagram of the sampling process conformations of Rosetta- distance, MMpred-C and MMpred and the comparison of the predicted first model with the native structure. The RMSD of the conformations sampled by Rosetta-distance are all greater than 3Å. Although the last generation conformations are diverse, the RMSD of most conformations is between 5 and12 Å. The final model clustered by SPICKER is 5.86Å. The RMSD of the conformations sampled by MMpred-C are all greater than 5 Å, and the conformations in the last generation population are concentrated. In contrast, MMpred can sample conformations with an RMSD less than 2 Å, and the conformations in the last generation of population are sampled in different basins. MMpred not only sampled the same low-energy basin (black arrow in **Figure 9C**) as that sampled by Rosetta-distance and MMpred-C, but also explored a more promising basin (yellow arrow in **Figure 9C**). The accuracy of the first model predicted by MMpred (RMSD = 2.93Å, TM-score = 0.80) is also significantly better than that of Rosetta-distance (RMSD = 5.86Å, TM-score = 0.61) and MMpred-C (RMSD = 5.95Å, TM-score = 0.56). This case shows that MMpred with multimodal optimization can effectively maintain the diversity of the population, explore more potential conformations of low-energy basins, and ultimately improve the accuracy of the prediction model.

**Fig. 9.**
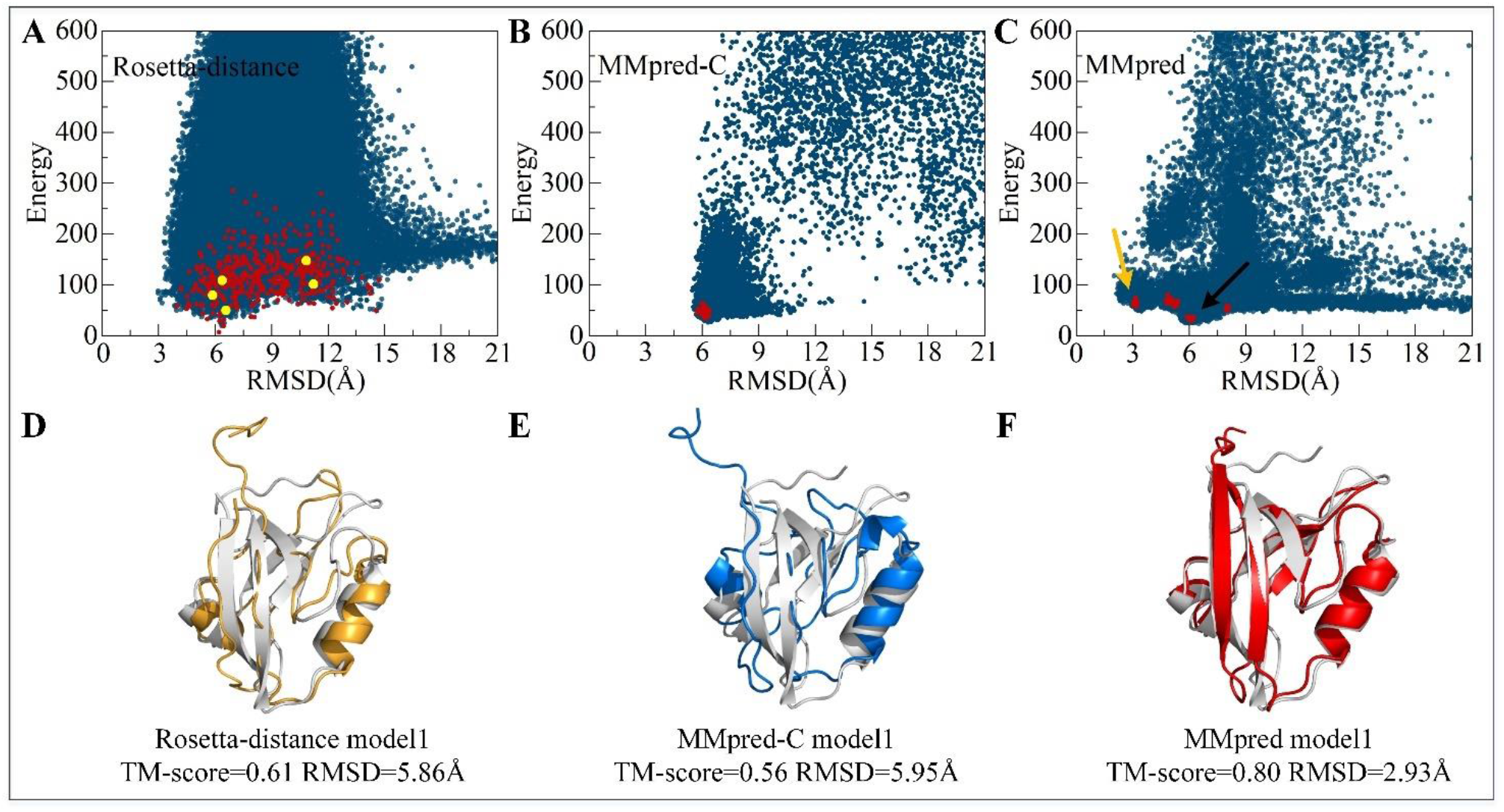
Example of comparing Rosetta-distance, MMpred-C and MMpred on the target 4Q2Q_A. (A), (B) and (C) are the RMSD-energy scatter diagram of process conformations. The process conformations is marked in blue, the last generation conformations is marked in red. The yellow points in (A) are the final five models of Rosetta-distance determined by SPICKER clustering. (D), (E) and (F) show the comparison of the predicted first model with the native structure (gray).

## 4 Conclusion

A distance-assisted multimodal optimization sampling algorithm, MMpred, is proposed. MMpred consists of three stages. In the first modal exploration stage, a structural similarity evaluation model DMscore is designed to control the diversity of conformations and generate the population with structural diversity. In the second modal maintaining stage, an adaptive clustering algorithm MNDcluster is proposed to divide the population, and then merge the modal by adjusting the annealing temperature. In the final modal exploitation stage, a greedy search strategy is used to accelerate the convergence of the modal. In addition, a distance scoring model based on the predicted distance constraint is used to guide conformational sampling together with the energy function. MMpred was compared with Rosetta-distance using distance constraint on 320 benchmark proteins. MMpred obtains models with TM-score?0.5 on 268 benchmark proteins, accounting for 84% of the total, which is better than Rosetta-distance. The comparison of MMpred and Rosetta-distance reflects the contribution of multimodal optimization to improving prediction accuracy. Furthermore, MMpred is tested on 24 FM targets of CASP 13 and the result is comparable with the four state-of-the-art CASP 13 server groups.

It is important to note that the multimodal optimization strategy in MMpred is an independent protocol proposed for improving conformational sampling of protein folding simulations. Although as an illustrative example, it has been applied to the Rosetta platform in this study, the MMpred protocol can be used for improving the sampling and model accuracy on other advanced folding simulation programs, such as C-QUARK, or combine the modeling advantages of both platforms of Rosetta and C-QUARK. In addition, combining molecular dynamics and intelligent optimization is expected to further refine model. Integrating the local abstract convex underestimation (Zhou and Zhang, 2017; Zhou et al., 2016) into the modal exploration is a good direction to improve the efficiency of the algorithm. Studies along these lines is under progress.

## Supplementary Information

**Figure S1:**
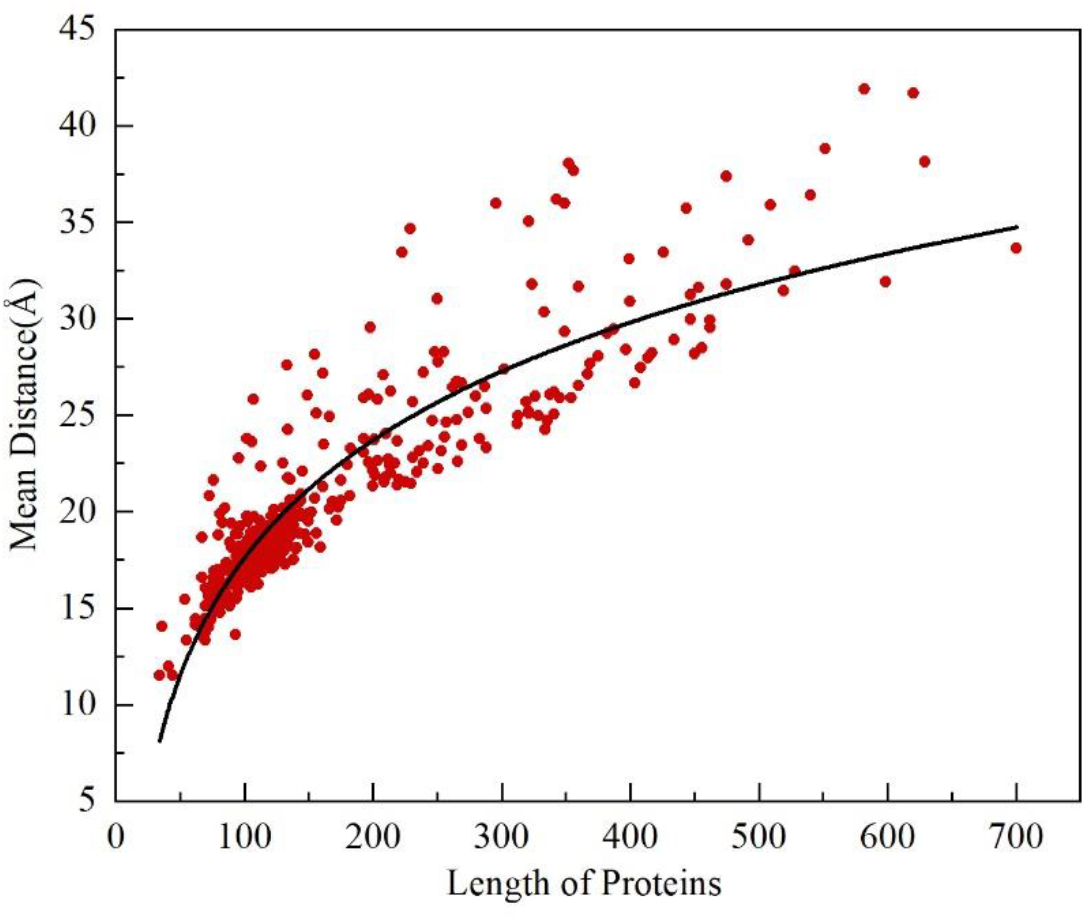
The relationship between the protein size and the mean Euclidean distance of inter-residues of the protein structure. We experimented with 500 native structures with a length of 40 to 700. The average distance between residues increases logarithmically as the length of the protein increases.

**Figure S2:**
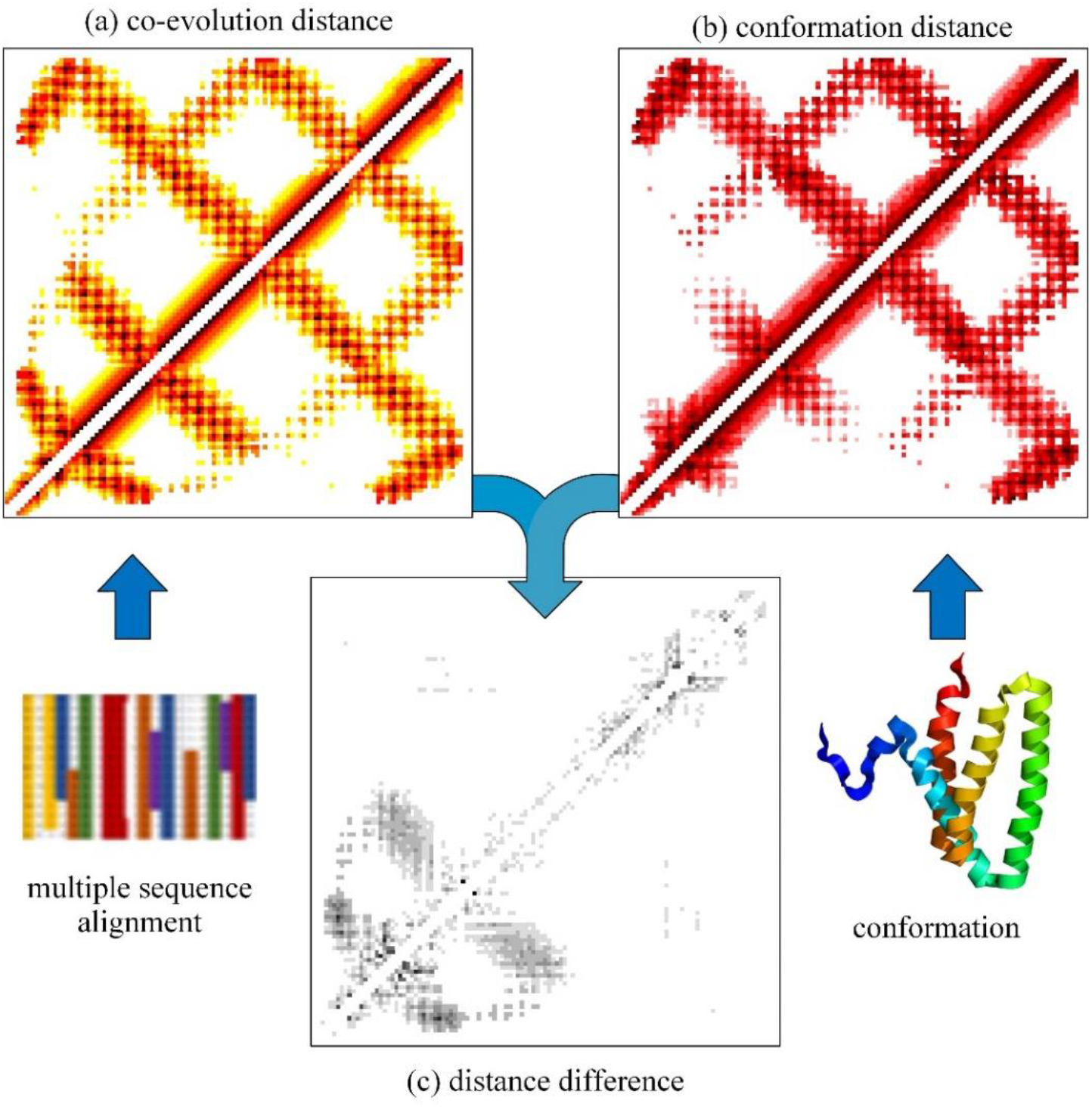
(a) is the inter-residue distance map predicted, (b) is the inter-residue distance map of sampled conformation, and (c) is the distance difference map after the superposition of (a) and (b).

**Figure S3:**
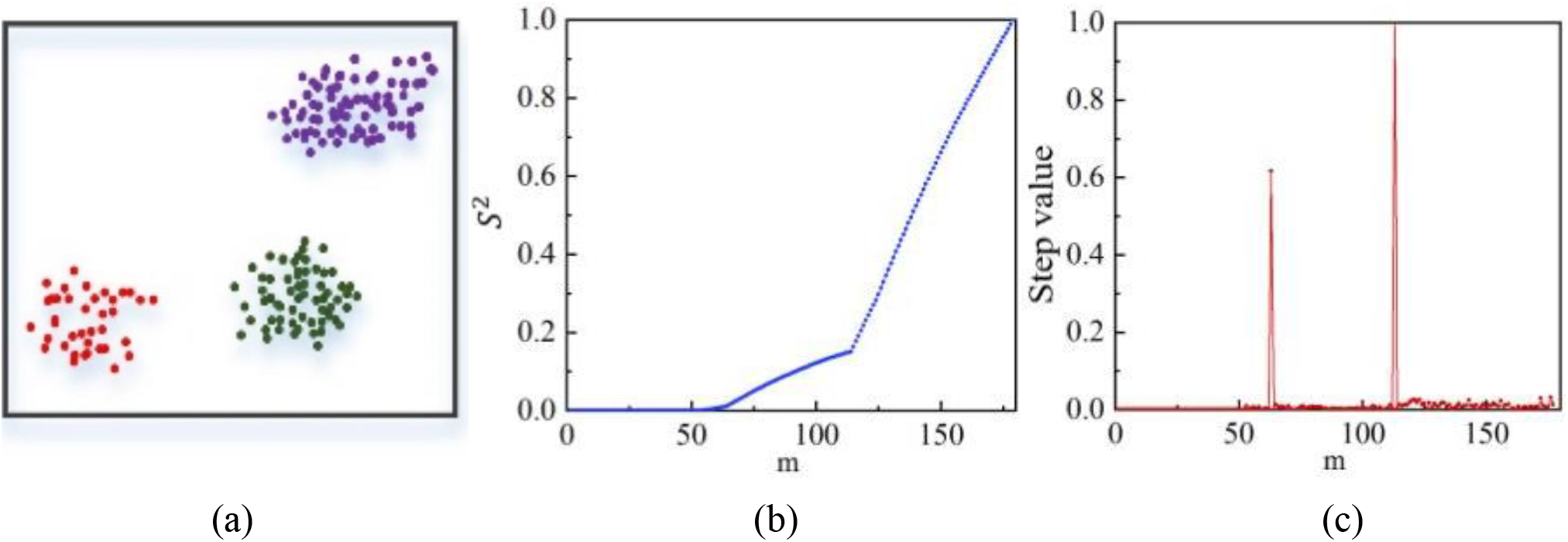
Schematic of MNDcluster. (a) Population distribution. (b) The relationship between the number of nearest neighbours and the variance. (c) Slope change of nearest neighbours values.

**Figure S4:**
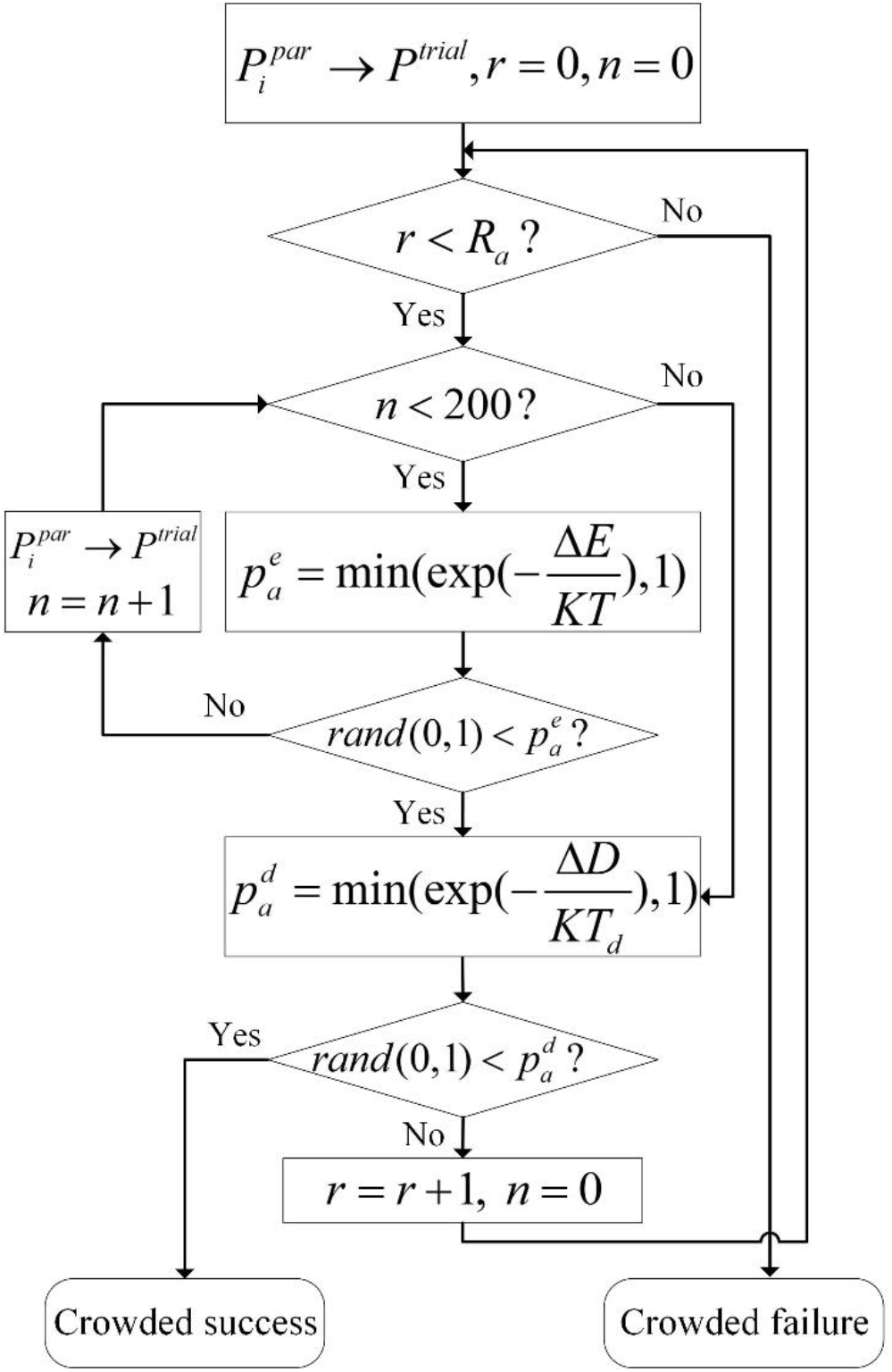
Flowchart of crowding fragment assembly in modal exploration stage. Where Δ*E* = *E^trial^ – E^par^*, Δ*D* = *D^trial^ -D^similar^*, *D^similar^* is the distance score of the conformation with the most similar conformation to trial conformation in the population; *R_a_* is crowding subpopulation size; *KT* and *KT_d_* are the temperature scaling factor. Trial conformation are produced by piecework assembly, The conformation is received according to the 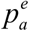 based on the energy score. If received fails, fragment assembly is performed again, up to 200 times. And then conformation is crowded according to the 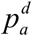 based on the distance score. Perform the iterating operation in sequence with the top *R* most similar conformations until the crowding is successful.

**Figure S5:**
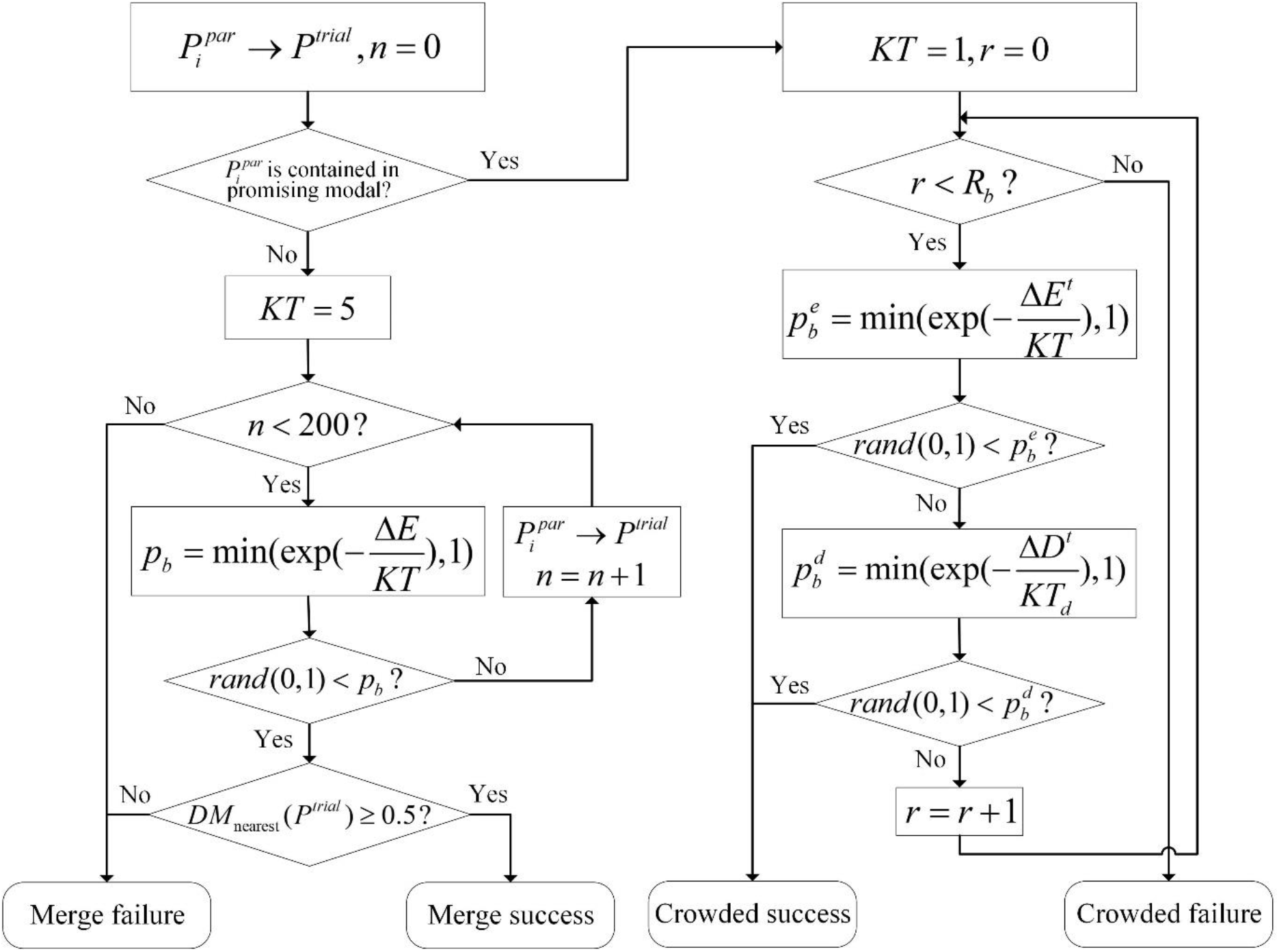
Flowchart of conformation update strategy in modal maintaining stage. Where Δ*E* = *E^trial^ – E^par^*, Δ*E^t^* = *E^trial^ – E^similar^*, Δ*D* = *D^trial^ – D^similar^*, *D^similar^* is the distance score of the conformation with the most similar conformation to trial conformation in the population; *R_b_* is crowding subpopulation size; If the conformation contained in the promising modal, the temperature is raised. After successful acceptance according to *P_b_*, the DMscore between the trial conformation of the unpromising modal and the centroid of the nearest promising modal was calculated. When DMscore is greater than or equal to 0.5, the conformations will merge into the promising modal. If the conformation contained in the unpromising modal, then the temperature is reduced. Metropolis criterion is used to determine whether the target conformation is crowded by trial conformation.

**Figure S6:**
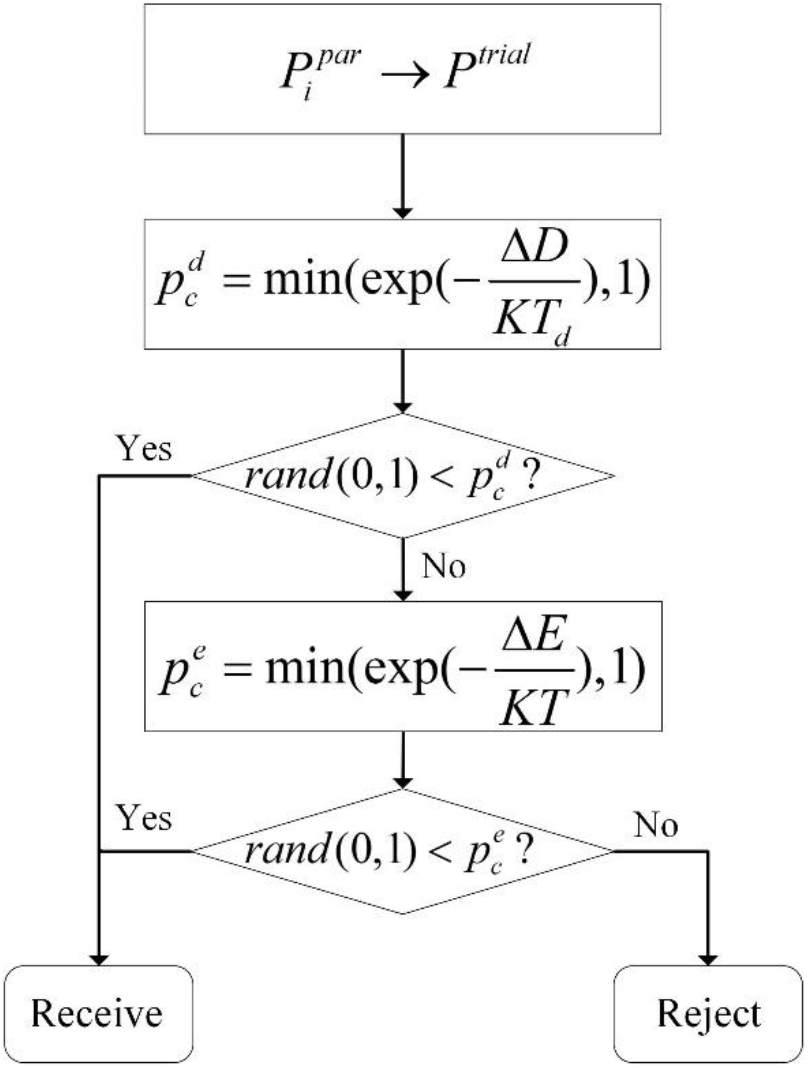
Flowchart of greedy fragment assembly in modal exploitation stage. Where Δ*D* = *D^trial^ – D^min^*, Δ*E* = *E^trial^ – E^min^*, *D^min^* is the lowest distance score and *E^min^* is the lowest energy score in the population.

**Figure S7:**
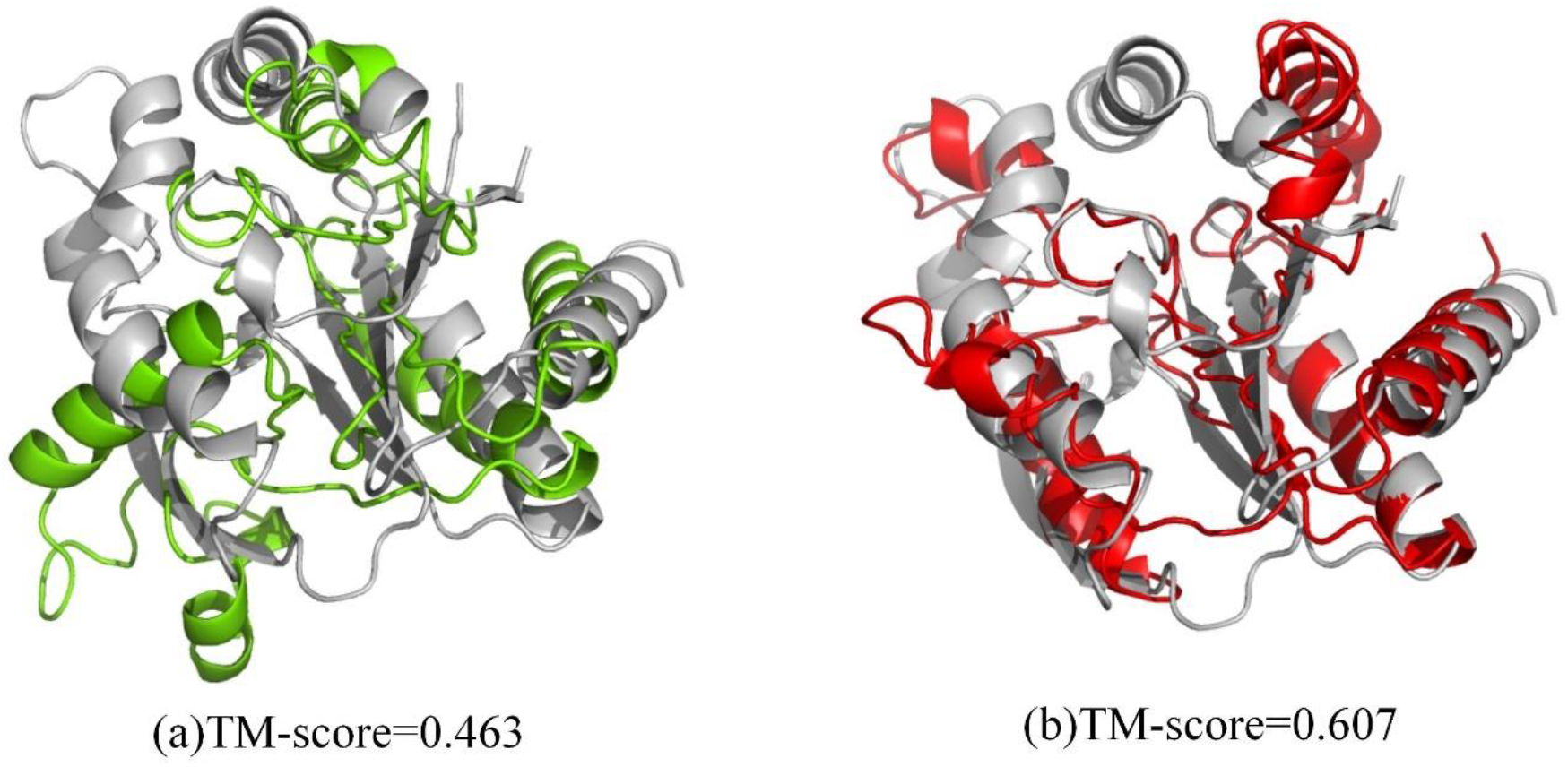
(a) is the predicted result of MMpred with a population size of 400 and generation of modal exploitation of 800. (b) is the predicted result of MMpred with a population size of 800 and generation of modal exploitation of 2000.

**Figure S8:**
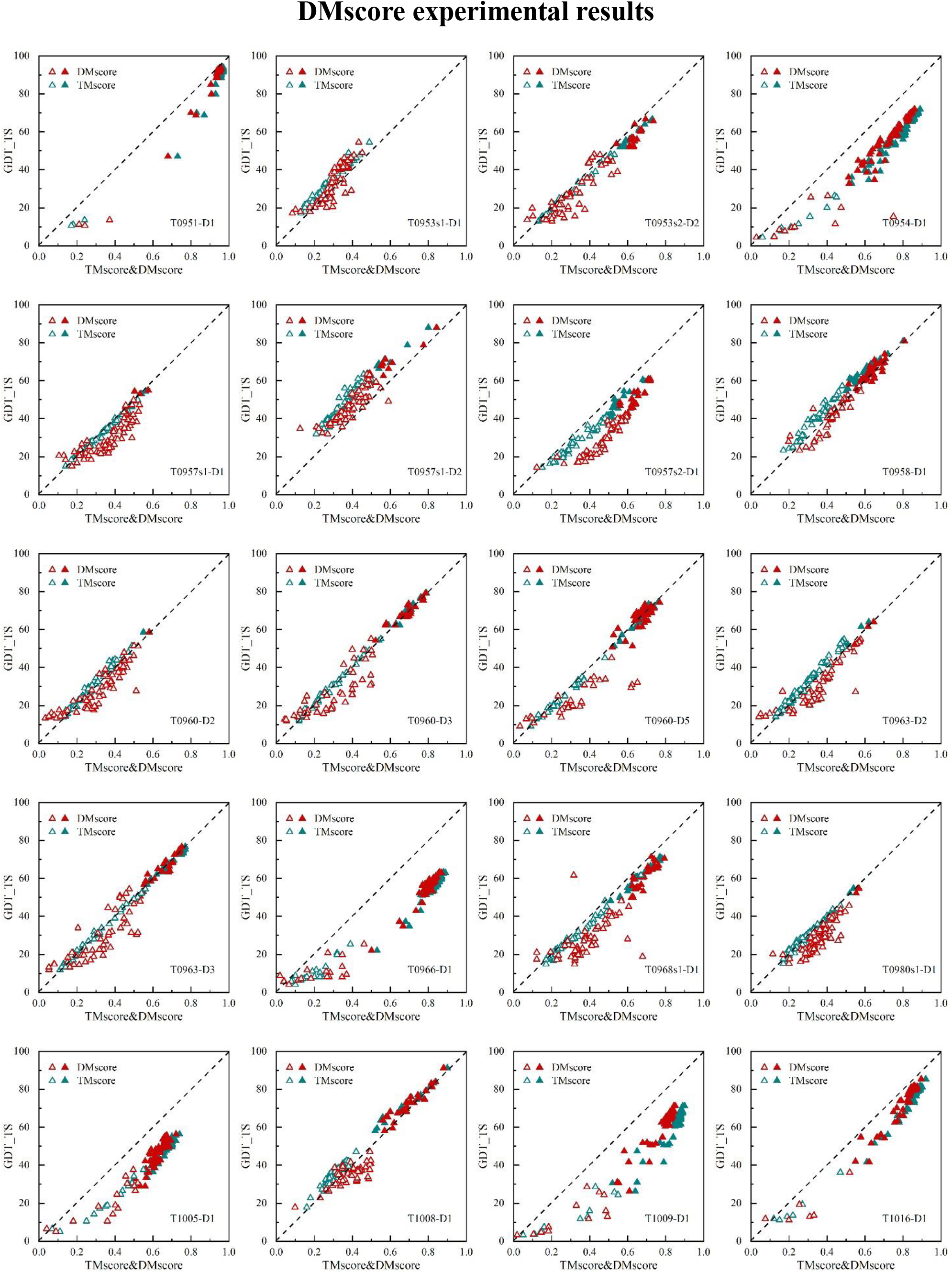
The relationship between GDT_TS and DMscore/TM-score. DMscore tested on 20 targets of CASP13, each containing the first model predicted by all the groups. Red triangle represents DMscore, blue triangle represents TM-score; Solid triangle means that DMscore and TM-score are both greater than or equal to 0.5.

**Table S1:** Prediction results (TM-score) of MMpred, Rosetta-distance and C-QUARK for 320 benchmark proteins.

| No. | PDB | Type | Size | MMPred |  | Rosetta-distance |  | C-QUARK |  |
| --- | --- | --- | --- | --- | --- | --- | --- | --- | --- |
|  |  |  |  | First model | Best model | First model | Best model | First model | Best model |
| 1 | 1FAQ_A | $\beta$ | 52 | 0.596 | 0.634 | 0.185 | 0.273 | 0.402 | 0.425 |
| 2 | 1JEI_A | $\alpha$ | 53 | 0.628 | 0.637 | 0.581 | 0.581 | 0.610 | 0.619 |
| 3 | 1NKZ_A | $\alpha$ | 53 | 0.571 | 0.587 | 0.289 | 0.357 | 0.417 | 0.424 |
| 4 | 1AUU_A | $\beta$ | 55 | 0.599 | 0.652 | 0.437 | 0.437 | 0.692 | 0.692 |
| 5 | 1FCA_A | $\alpha/\beta$ | 55 | 0.814 | 0.845 | 0.690 | 0.753 | 0.675 | 0.675 |
| 6 | 2UUX_A | $\alpha/\beta$ | 55 | 0.678 | 0.682 | 0.186 | 0.267 | 0.355 | 0.464 |
| 7 | 1H9E_A | $\alpha$ | 56 | 0.504 | 0.536 | 0.492 | 0.492 | 0.521 | 0.568 |
| 8 | 1H9F_A | $\alpha$ | 57 | 0.565 | 0.568 | 0.542 | 0.542 | 0.595 | 0.597 |
| 9 | 1LR1_B | $\alpha$ | 57 | 0.498 | 0.507 | 0.482 | 0.520 | 0.434 | 0.437 |
| 10 | 1DL6_A | $\beta$ | 58 | 0.392 | 0.392 | 0.383 | 0.411 | 0.369 | 0.374 |
| 11 | 1E53_A | $\alpha/\beta$ | 59 | 0.620 | 0.620 | 0.470 | 0.547 | 0.516 | 0.516 |
| 12 | 1FEX_A | $\alpha$ | 59 | 0.645 | 0.662 | 0.568 | 0.568 | 0.587 | 0.592 |
| 13 | 1UG4_A | $\beta$ | 60 | 0.589 | 0.625 | 0.320 | 0.320 | 0.440 | 0.440 |
| 14 | 1VYX_A | $\alpha/\beta$ | 60 | 0.444 | 0.486 | 0.270 | 0.270 | 0.433 | 0.450 |
| 15 | 3P8B_A | $\alpha/\beta$ | 60 | 0.518 | 0.666 | 0.296 | 0.296 | 0.669 | 0.669 |
| 16 | 1F43_A | $\alpha$ | 61 | 0.563 | 0.576 | 0.560 | 0.587 | 0.559 | 0.576 |
| 17 | 1L2P_A | $\alpha$ | 61 | 0.864 | 0.939 | 0.300 | 0.326 | 0.458 | 0.490 |
| 18 | 4MLF_D | $\beta$ | 61 | 0.284 | 0.284 | 0.206 | 0.226 | 0.247 | 0.248 |
| 19 | 1BE3_J | $\alpha$ | 62 | 0.544 | 0.605 | 0.365 | 0.365 | 0.271 | 0.287 |
| 20 | 1BGY_J | $\alpha$ | 62 | 0.540 | 0.567 | 0.400 | 0.400 | 0.270 | 0.292 |
| 21 | 1TYG_B | $\alpha/\beta$ | 65 | 0.797 | 0.823 | 0.632 | 0.648 | 0.803 | 0.818 |
| 22 | 1DTV_A | $\alpha/\beta$ | 67 | 0.226 | 0.255 | 0.204 | 0.204 | 0.362 | 0.428 |
| 23 | 2JP3_A | $\alpha$ | 67 | 0.364 | 0.389 | 0.342 | 0.348 | 0.257 | 0.257 |
| 24 | 1CHC_A | $\alpha/\beta$ | 68 | 0.489 | 0.501 | 0.290 | 0.381 | 0.504 | 0.545 |
| 25 | 1J9I_A | $\alpha/\beta$ | 68 | 0.623 | 0.623 | 0.462 | 0.493 | 0.632 | 0.655 |
| 26 | 1TAF_A | $\alpha$ | 68 | 0.859 | 0.874 | 0.533 | 0.576 | 0.513 | 0.530 |
| 27 | 1DWM_A | $\alpha/\beta$ | 69 | 0.688 | 0.688 | 0.565 | 0.565 | 0.643 | 0.653 |
| 28 | 1L6H_A | $\alpha$ | 69 | 0.516 | 0.519 | 0.508 | 0.508 | 0.413 | 0.416 |
| 29 | 1W1W_E | $\alpha/\beta$ | 70 | 0.805 | 0.834 | 0.807 | 0.807 | 0.813 | 0.813 |
| 30 | 2CMX_A | $\alpha/\beta$ | 70 | 0.709 | 0.760 | 0.605 | 0.638 | 0.596 | 0.704 |
| 31 | 1LDD_A | $\alpha/\beta$ | 71 | 0.850 | 0.862 | 0.354 | 0.669 | 0.708 | 0.732 |
| 32 | 1K73_1 | $\alpha/\beta$ | 73 | 0.642 | 0.682 | 0.468 | 0.468 | 0.447 | 0.469 |
| 33 | 1KN6_A | $\alpha/\beta$ | 73 | 0.575 | 0.594 | 0.512 | 0.512 | 0.591 | 0.591 |
| 34 | 1PIH_A | $\alpha/\beta$ | 73 | 0.558 | 0.604 | 0.288 | 0.288 | 0.519 | 0.564 |
| 35 | 2HI3_A | $\alpha$ | 73 | 0.641 | 0.643 | 0.654 | 0.673 | 0.695 | 0.704 |
| 36 | 1ABT_A | $\alpha/\beta$ | 74 | 0.484 | 0.484 | 0.373 | 0.417 | 0.496 | 0.496 |
| 37 | 1HHV_A | $\alpha/\beta$ | 74 | 0.598 | 0.605 | 0.439 | 0.439 | 0.628 | 0.628 |
| 38 | 2V85_A | $\alpha/\beta$ | 74 | 0.430 | 0.464 | 0.284 | 0.325 | 0.275 | 0.346 |
| 39 | 3CX5_F | $\alpha$ | 74 | 0.788 | 0.788 | 0.501 | 0.517 | 0.709 | 0.735 |
| 40 | 3MQK_C | $\beta$ | 75 | 0.805 | 0.850 | 0.539 | 0.539 | 0.754 | 0.756 |
| 41 | 4M75_F | $\alpha/\beta$ | 75 | 0.742 | 0.742 | 0.574 | 0.574 | 0.719 | 0.724 |
| 42 | 1DP7_P | $\alpha/\beta$ | 76 | 0.745 | 0.770 | 0.489 | 0.531 | 0.744 | 0.748 |
| 43 | 1EKZ_A | $\alpha/\beta$ | 76 | 0.674 | 0.693 | 0.591 | 0.664 | 0.690 | 0.693 |
| 44 | 1VCC_A | $\alpha/\beta$ | 77 | 0.557 | 0.559 | 0.517 | 0.517 | 0.552 | 0.594 |
| 45 | 1K1Z_A | $\beta$ | 78 | 0.519 | 0.541 | 0.503 | 0.503 | 0.602 | 0.624 |
| 46 | 1PFS_A | $\beta$ | 78 | 0.530 | 0.547 | 0.327 | 0.383 | 0.538 | 0.567 |
| 47 | 4V2O_A | $\alpha$ | 78 | 0.773 | 0.804 | 0.592 | 0.592 | 0.545 | 0.545 |
| 48 | 1A91_A | $\alpha$ | 79 | 0.506 | 0.540 | 0.562 | 0.562 | 0.475 | 0.503 |
| 49 | 1KP6_A | $\alpha/\beta$ | 79 | 0.242 | 0.280 | 0.293 | 0.293 | 0.330 | 0.351 |
| 50 | 2ZMZ_B | $\alpha/\beta$ | 79 | 0.713 | 0.729 | 0.497 | 0.563 | 0.456 | 0.611 |

| No. | PDB | Type | Size | MMpred |  | Rosetta-distance |  | C-QUARK |  |
| --- | --- | --- | --- | --- | --- | --- | --- | --- | --- |
|  |  |  |  | First model | Best model | First model | Best model | First model | Best model |
| 51 | 1B4R_A | $\beta$ | 80 | 0.650 | 0.717 | 0.553 | 0.553 | 0.718 | 0.756 |
| 52 | 1PZW_A | $\alpha/\beta$ | 80 | 0.717 | 0.762 | 0.287 | 0.335 | 0.523 | 0.533 |
| 53 | 1R6R_A | $\alpha$ | 80 | 0.531 | 0.531 | 0.323 | 0.406 | 0.385 | 0.397 |
| 54 | 4LMS_A | $\alpha/\beta$ | 80 | 0.402 | 0.402 | 0.269 | 0.269 | 0.346 | 0.346 |
| 55 | 1D8B_A | $\alpha$ | 81 | 0.732 | 0.744 | 0.684 | 0.684 | 0.659 | 0.690 |
| 56 | 1EZV_G | $\alpha/\beta$ | 81 | 0.374 | 0.418 | 0.371 | 0.397 | 0.324 | 0.324 |
| 57 | 1F9P_A | $\alpha/\beta$ | 81 | 0.593 | 0.599 | 0.548 | 0.569 | 0.611 | 0.629 |
| 58 | 1GGS_A | $\alpha/\beta$ | 81 | 0.681 | 0.695 | 0.589 | 0.625 | 0.659 | 0.679 |
| 59 | 1U84_A | $\alpha$ | 81 | 0.840 | 0.848 | 0.633 | 0.701 | 0.800 | 0.828 |
| 60 | 4ASW_C | $\alpha/\beta$ | 81 | 0.726 | 0.778 | 0.534 | 0.534 | 0.710 | 0.721 |
| 61 | 1CF7_B | $\alpha/\beta$ | 82 | 0.718 | 0.776 | 0.520 | 0.520 | 0.684 | 0.689 |
| 62 | 1LFU_P | $\alpha$ | 82 | 0.589 | 0.590 | 0.593 | 0.613 | 0.567 | 0.567 |
| 63 | 1WMH_B | $\alpha/\beta$ | 82 | 0.782 | 0.795 | 0.459 | 0.495 | 0.733 | 0.783 |
| 64 | 2ODM_B | $\alpha$ | 83 | 0.826 | 0.832 | 0.780 | 0.795 | 0.682 | 0.731 |
| 65 | 2AQ0_A | $\alpha$ | 84 | 0.644 | 0.651 | 0.627 | 0.646 | 0.630 | 0.644 |
| 66 | 2GBJ_B | $\alpha/\beta$ | 84 | 0.749 | 0.754 | 0.551 | 0.623 | 0.779 | 0.804 |
| 67 | 1IS7_K | $\alpha/\beta$ | 85 | 0.643 | 0.643 | 0.417 | 0.417 | 0.484 | 0.484 |
| 68 | 5T17_A | $\alpha/\beta$ | 85 | 0.753 | 0.753 | 0.729 | 0.729 | 0.724 | 0.753 |
| 69 | 1CXZ_B | $\alpha$ | 86 | 0.867 | 0.886 | 0.708 | 0.708 | 0.797 | 0.850 |
| 70 | 1NOE_A | $\alpha/\beta$ | 86 | 0.611 | 0.611 | 0.271 | 0.302 | 0.515 | 0.515 |
| 71 | 2J6Z_A | $\alpha$ | 86 | 0.720 | 0.739 | 0.413 | 0.586 | 0.666 | 0.674 |
| 72 | 4OW1_A | $\alpha/\beta$ | 86 | 0.738 | 0.741 | 0.564 | 0.564 | 0.657 | 0.712 |
| 73 | 1C9F_A | $\alpha/\beta$ | 87 | 0.578 | 0.589 | 0.552 | 0.572 | 0.601 | 0.616 |
| 74 | 1GVP_A | $\alpha/\beta$ | 87 | 0.671 | 0.671 | 0.479 | 0.479 | 0.595 | 0.595 |
| 75 | 1RHX_A | $\alpha/\beta$ | 87 | 0.580 | 0.595 | 0.511 | 0.511 | 0.568 | 0.584 |
| 76 | 3X15_A | $\alpha$ | 87 | 0.498 | 0.498 | 0.261 | 0.261 | 0.445 | 0.458 |
| 77 | 4GDK_A | $\alpha/\beta$ | 88 | 0.671 | 0.694 | 0.505 | 0.505 | 0.696 | 0.746 |
| 78 | 4J20_A | $\alpha/\beta$ | 88 | 0.704 | 0.743 | 0.558 | 0.558 | 0.553 | 0.592 |
| 79 | 1H8E_H | $\alpha/\beta$ | 89 | 0.657 | 0.657 | 0.519 | 0.519 | 0.775 | 0.809 |
| 80 | 1HBX_E | $\alpha/\beta$ | 89 | 0.516 | 0.527 | 0.523 | 0.523 | 0.414 | 0.451 |
| 81 | 5IZB_A | $\alpha/\beta$ | 89 | 0.565 | 0.565 | 0.441 | 0.441 | 0.557 | 0.584 |
| 82 | 1JR5_A | $\alpha$ | 90 | 0.522 | 0.572 | 0.349 | 0.446 | 0.345 | 0.461 |
| 83 | 4Q2Q_A | $\alpha/\beta$ | 90 | 0.801 | 0.801 | 0.612 | 0.612 | 0.703 | 0.803 |
| 84 | 4MMG_A | $\alpha/\beta$ | 91 | 0.823 | 0.840 | 0.633 | 0.633 | 0.727 | 0.763 |
| 85 | 5L38_A | $\alpha/\beta$ | 91 | 0.825 | 0.843 | 0.664 | 0.667 | 0.856 | 0.856 |
| 86 | 1IUY_A | $\alpha/\beta$ | 92 | 0.689 | 0.697 | 0.458 | 0.458 | 0.677 | 0.677 |
| 87 | 2BYK_D | $\alpha$ | 92 | 0.856 | 0.876 | 0.575 | 0.655 | 0.570 | 0.600 |
| 88 | 4Q2O_A | $\alpha/\beta$ | 92 | 0.665 | 0.665 | 0.615 | 0.615 | 0.776 | 0.798 |
| 89 | 5O2V_A | $\alpha/\beta$ | 92 | 0.747 | 0.747 | 0.590 | 0.626 | 0.775 | 0.794 |
| 90 | 1J8I_A | $\alpha/\beta$ | 93 | 0.538 | 0.555 | 0.539 | 0.571 | 0.568 | 0.579 |
| 91 | 3X0G_A | $\alpha$ | 93 | 0.533 | 0.533 | 0.322 | 0.445 | 0.559 | 0.603 |
| 92 | 1G2R_A | $\alpha/\beta$ | 94 | 0.811 | 0.819 | 0.655 | 0.655 | 0.771 | 0.787 |
| 93 | 1PD6_A | $\beta$ | 94 | 0.598 | 0.634 | 0.607 | 0.607 | 0.662 | 0.662 |
| 94 | 2ICT_A | $\alpha$ | 94 | 0.744 | 0.763 | 0.707 | 0.712 | 0.750 | 0.765 |
| 95 | 1I35_A | $\alpha/\beta$ | 95 | 0.573 | 0.573 | 0.457 | 0.457 | 0.439 | 0.482 |
| 96 | 1IM3_D | $\alpha/\beta$ | 95 | 0.264 | 0.328 | 0.219 | 0.298 | 0.278 | 0.282 |
| 97 | 5TMF_E | $\alpha/\beta$ | 95 | 0.553 | 0.567 | 0.388 | 0.388 | 0.505 | 0.505 |
| 98 | 3N9U_C | $\alpha/\beta$ | 96 | 0.731 | 0.736 | 0.586 | 0.640 | 0.758 | 0.770 |
| 99 | 1JO0_A | $\alpha/\beta$ | 97 | 0.820 | 0.852 | 0.415 | 0.599 | 0.806 | 0.839 |
| 100 | 2LWP_A | $\alpha/\beta$ | 97 | 0.501 | 0.557 | 0.458 | 0.486 | 0.541 | 0.552 |
|  |  |  |  | First model | Best model | First model | Best model | First model | Best model |
| 101 | 3SDL_B | $\alpha$ | 97 | 0.315 | 0.329 | 0.351 | 0.351 | 0.295 | 0.327 |
| 102 | 1FRD_A | $\alpha/\beta$ | 98 | 0.705 | 0.727 | 0.565 | 0.572 | 0.704 | 0.785 |
| 103 | 1JMT_A | $\alpha/\beta$ | 98 | 0.752 | 0.752 | 0.578 | 0.578 | 0.701 | 0.716 |
| 104 | 2ACY_A | $\alpha/\beta$ | 98 | 0.657 | 0.670 | 0.589 | 0.599 | 0.859 | 0.863 |
| 105 | 1MWQ_A | $\alpha/\beta$ | 99 | 0.674 | 0.674 | 0.599 | 0.599 | 0.576 | 0.583 |
| 106 | 2NCM_A | $\beta$ | 99 | 0.834 | 0.834 | 0.616 | 0.616 | 0.861 | 0.861 |
| 107 | 2NDP_A | $\alpha/\beta$ | 99 | 0.480 | 0.520 | 0.441 | 0.477 | 0.416 | 0.416 |
| 108 | 1F2R_I | $\alpha/\beta$ | 100 | 0.627 | 0.643 | 0.431 | 0.549 | 0.643 | 0.646 |
| 109 | 1KA8_A | $\alpha/\beta$ | 100 | 0.560 | 0.601 | 0.340 | 0.505 | 0.470 | 0.504 |
| 110 | 1PSR_A | $\alpha/\beta$ | 100 | 0.752 | 0.752 | 0.604 | 0.604 | 0.660 | 0.733 |
| 111 | 1SMP_I | $\alpha/\beta$ | 100 | 0.700 | 0.700 | 0.340 | 0.411 | 0.694 | 0.694 |
| 112 | 4IOS_A | $\alpha/\beta$ | 100 | 0.549 | 0.593 | 0.431 | 0.431 | 0.460 | 0.558 |
| 113 | 1FJG_F | $\alpha/\beta$ | 101 | 0.721 | 0.726 | 0.618 | 0.618 | 0.809 | 0.809 |
| 114 | 1TUL_A | $\alpha/\beta$ | 102 | 0.378 | 0.505 | 0.260 | 0.328 | 0.460 | 0.528 |
| 115 | 2K9X_A | $\alpha/\beta$ | 102 | 0.536 | 0.536 | 0.487 | 0.487 | 0.621 | 0.655 |
| 116 | 1F93_A | $\alpha/\beta$ | 103 | 0.779 | 0.790 | 0.681 | 0.689 | 0.860 | 0.860 |
| 117 | 1IUJ_B | $\alpha/\beta$ | 103 | 0.730 | 0.762 | 0.706 | 0.708 | 0.706 | 0.736 |
| 118 | 4ESB_A | $\alpha/\beta$ | 103 | 0.844 | 0.844 | 0.762 | 0.762 | 0.687 | 0.704 |
| 119 | 1E3Y_A | $\alpha$ | 104 | 0.713 | 0.731 | 0.618 | 0.618 | 0.685 | 0.690 |
| 120 | 1FMB_A | $\alpha/\beta$ | 104 | 0.528 | 0.599 | 0.352 | 0.402 | 0.633 | 0.637 |
| 121 | 3N1G_C | $\alpha/\beta$ | 104 | 0.642 | 0.658 | 0.620 | 0.627 | 0.818 | 0.818 |
| 122 | 4UIJ_A | $\alpha/\beta$ | 104 | 0.759 | 0.795 | 0.618 | 0.618 | 0.766 | 0.788 |
| 123 | 5TUV_B | $\alpha/\beta$ | 104 | 0.343 | 0.394 | 0.423 | 0.423 | 0.388 | 0.388 |
| 124 | 1ABV_A | $\alpha$ | 105 | 0.795 | 0.812 | 0.755 | 0.755 | 0.758 | 0.794 |
| 125 | 1CDB_A | $\beta$ | 105 | 0.625 | 0.625 | 0.541 | 0.541 | 0.665 | 0.665 |
| 126 | 1JIW_I | $\alpha/\beta$ | 105 | 0.539 | 0.539 | 0.462 | 0.464 | 0.662 | 0.662 |
| 127 | 1KPT_A | $\alpha/\beta$ | 105 | 0.623 | 0.667 | 0.326 | 0.326 | 0.619 | 0.638 |
| 128 | 1A6L_A | $\alpha/\beta$ | 106 | 0.658 | 0.684 | 0.503 | 0.503 | 0.360 | 0.520 |
| 129 | 1ID2_A | $\alpha/\beta$ | 106 | 0.566 | 0.679 | 0.499 | 0.499 | 0.656 | 0.675 |
| 130 | 1S7Z_A | $\alpha$ | 106 | 0.554 | 0.588 | 0.576 | 0.576 | 0.686 | 0.736 |
| 131 | 4RUV_A | $\alpha/\beta$ | 106 | 0.857 | 0.857 | 0.832 | 0.844 | 0.850 | 0.854 |
| 132 | 1KX5_D | $\alpha$ | 107 | 0.708 | 0.775 | 0.524 | 0.530 | 0.477 | 0.497 |
| 133 | 1ROW_A | $\alpha/\beta$ | 107 | 0.549 | 0.600 | 0.471 | 0.471 | 0.677 | 0.688 |
| 134 | 1V74_A | $\alpha/\beta$ | 107 | 0.727 | 0.727 | 0.539 | 0.540 | 0.596 | 0.644 |
| 135 | 2BSE_A | $\alpha/\beta$ | 107 | 0.448 | 0.462 | 0.295 | 0.329 | 0.306 | 0.344 |
| 136 | 1MFQ_C | $\alpha$ | 108 | 0.637 | 0.644 | 0.608 | 0.611 | 0.583 | 0.603 |
| 137 | 1S7O_C | $\alpha$ | 108 | 0.574 | 0.574 | 0.511 | 0.644 | 0.679 | 0.679 |
| 138 | 1TZ0_A | $\alpha/\beta$ | 108 | 0.545 | 0.618 | 0.527 | 0.527 | 0.650 | 0.658 |
| 139 | 1K3S_A | $\alpha/\beta$ | 109 | 0.616 | 0.690 | 0.496 | 0.518 | 0.299 | 0.453 |
| 140 | 1VD0_A | $\alpha/\beta$ | 109 | 0.433 | 0.479 | 0.220 | 0.303 | 0.599 | 0.610 |
| 141 | 2CWP_A | $\alpha/\beta$ | 109 | 0.659 | 0.736 | 0.507 | 0.507 | 0.725 | 0.725 |
| 142 | 2NAZ_A | $\alpha/\beta$ | 109 | 0.661 | 0.698 | 0.618 | 0.654 | 0.747 | 0.747 |
| 143 | 1BJX_A | $\alpha/\beta$ | 110 | 0.786 | 0.786 | 0.598 | 0.599 | 0.741 | 0.785 |
| 144 | 1I85_A | $\beta$ | 110 | 0.598 | 0.628 | 0.641 | 0.641 | 0.605 | 0.632 |
| 145 | 1QFW_B | $\beta$ | 110 | 0.383 | 0.383 | 0.330 | 0.332 | 0.372 | 0.395 |
| 146 | 2D0P_B | $\alpha/\beta$ | 110 | 0.683 | 0.752 | 0.659 | 0.659 | 0.747 | 0.747 |
| 147 | 3W1Z_D | $\alpha/\beta$ | 110 | 0.569 | 0.624 | 0.530 | 0.554 | 0.597 | 0.636 |
| 148 | 1EM8_D | $\alpha/\beta$ | 112 | 0.649 | 0.712 | 0.499 | 0.499 | 0.568 | 0.579 |
| 149 | 1JLI_A | $\alpha$ | 112 | 0.365 | 0.378 | 0.205 | 0.260 | 0.336 | 0.466 |
| 150 | 1NZE_A | $\alpha$ | 112 | 0.867 | 0.878 | 0.826 | 0.826 | 0.869 | 0.875 |
|  |  |  |  | First model | Best model | First model | Best model | First model | Best model |
| 151 | 1N3G_A | $\alpha/\beta$ | 113 | 0.692 | 0.699 | 0.654 | 0.654 | 0.753 | 0.753 |
| 152 | 1SAU_A | $\alpha/\beta$ | 114 | 0.678 | 0.705 | 0.355 | 0.475 | 0.672 | 0.787 |
| 153 | 2APN_A | $\alpha/\beta$ | 114 | 0.615 | 0.615 | 0.461 | 0.501 | 0.666 | 0.666 |
| 154 | 5WSE_A | $\alpha/\beta$ | 114 | 0.654 | 0.711 | 0.636 | 0.636 | 0.667 | 0.688 |
| 155 | 3EOD_A | $\alpha/\beta$ | 115 | 0.817 | 0.820 | 0.681 | 0.727 | 0.795 | 0.827 |
| 156 | 3LQV_B | $\alpha/\beta$ | 115 | 0.640 | 0.645 | 0.498 | 0.508 | 0.613 | 0.647 |
| 157 | 1FC3_A | $\alpha$ | 116 | 0.819 | 0.828 | 0.671 | 0.671 | 0.772 | 0.804 |
| 158 | 1FHT_A | $\alpha/\beta$ | 116 | 0.624 | 0.624 | 0.545 | 0.572 | 0.663 | 0.677 |
| 159 | 2RLD_C | $\alpha$ | 116 | 0.876 | 0.883 | 0.793 | 0.819 | 0.825 | 0.868 |
| 160 | 1D6T_A | $\alpha/\beta$ | 117 | 0.619 | 0.694 | 0.477 | 0.574 | 0.705 | 0.717 |
| 161 | 1ELW_A | $\alpha$ | 117 | 0.913 | 0.918 | 0.880 | 0.950 | 0.870 | 0.894 |
| 162 | 1WJ8_A | $\alpha$ | 117 | 0.872 | 0.875 | 0.758 | 0.758 | 0.839 | 0.861 |
| 163 | 1A7D_A | $\alpha$ | 118 | 0.789 | 0.791 | 0.641 | 0.641 | 0.760 | 0.791 |
| 164 | 1HCD_A | $\alpha/\beta$ | 118 | 0.626 | 0.626 | 0.323 | 0.372 | 0.671 | 0.724 |
| 165 | 2Q2H_A | $\alpha/\beta$ | 118 | 0.498 | 0.593 | 0.455 | 0.471 | 0.664 | 0.681 |
| 166 | 1JJ2_S | $\alpha/\beta$ | 119 | 0.669 | 0.727 | 0.435 | 0.435 | 0.437 | 0.453 |
| 167 | 1MAI_A | $\alpha/\beta$ | 119 | 0.611 | 0.703 | 0.535 | 0.535 | 0.705 | 0.708 |
| 168 | 1OFT_A | $\alpha/\beta$ | 119 | 0.799 | 0.799 | 0.704 | 0.704 | 0.776 | 0.776 |
| 169 | 1ORY_A | $\alpha$ | 119 | 0.783 | 0.783 | 0.722 | 0.722 | 0.731 | 0.736 |
| 170 | 1VKE_E | $\alpha$ | 119 | 0.790 | 0.790 | 0.619 | 0.645 | 0.642 | 0.642 |
| 171 | 2CZV_D | $\alpha/\beta$ | 119 | 0.681 | 0.710 | 0.372 | 0.378 | 0.670 | 0.734 |
| 172 | 2GJ3_A | $\alpha/\beta$ | 119 | 0.802 | 0.802 | 0.663 | 0.663 | 0.833 | 0.838 |
| 173 | 2LKP_A | $\alpha/\beta$ | 119 | 0.594 | 0.607 | 0.586 | 0.586 | 0.561 | 0.567 |
| 174 | 2PI2_F | $\alpha/\beta$ | 119 | 0.782 | 0.782 | 0.486 | 0.486 | 0.743 | 0.745 |
| 175 | 3G20_B | $\alpha/\beta$ | 119 | 0.570 | 0.570 | 0.454 | 0.454 | 0.618 | 0.618 |
| 176 | 1DUN_A | $\alpha/\beta$ | 120 | 0.609 | 0.609 | 0.419 | 0.447 | 0.764 | 0.786 |
| 177 | 1JPY_Y | $\alpha/\beta$ | 120 | 0.414 | 0.415 | 0.401 | 0.401 | 0.472 | 0.472 |
| 178 | 2H8E_A | $\alpha/\beta$ | 120 | 0.614 | 0.614 | 0.390 | 0.390 | 0.628 | 0.679 |
| 179 | 1BUO_A | $\alpha/\beta$ | 121 | 0.804 | 0.804 | 0.478 | 0.479 | 0.791 | 0.791 |
| 180 | 2QZJ_A | $\alpha/\beta$ | 121 | 0.895 | 0.895 | 0.800 | 0.852 | 0.923 | 0.923 |
| 181 | 1NR3_A | $\alpha/\beta$ | 122 | 0.316 | 0.362 | 0.292 | 0.361 | 0.272 | 0.294 |
| 182 | 2EWC_B | $\alpha/\beta$ | 122 | 0.684 | 0.701 | 0.594 | 0.615 | 0.746 | 0.760 |
| 183 | 2WCW_B | $\alpha/\beta$ | 122 | 0.714 | 0.759 | 0.634 | 0.634 | 0.749 | 0.759 |
| 184 | 3QU3_A | $\alpha/\beta$ | 122 | 0.552 | 0.552 | 0.472 | 0.472 | 0.508 | 0.587 |
| 185 | 5O8G_A | $\alpha/\beta$ | 122 | 0.641 | 0.710 | 0.534 | 0.563 | 0.772 | 0.772 |
| 186 | 1L3G_A | $\alpha/\beta$ | 123 | 0.531 | 0.583 | 0.513 | 0.538 | 0.557 | 0.557 |
| 187 | 1QMA_A | $\alpha/\beta$ | 123 | 0.653 | 0.749 | 0.640 | 0.640 | 0.893 | 0.893 |
| 188 | 4GF3_A | $\alpha/\beta$ | 123 | 0.577 | 0.637 | 0.574 | 0.574 | 0.658 | 0.658 |
| 189 | 1BGF_A | $\alpha/\beta$ | 124 | 0.801 | 0.816 | 0.362 | 0.449 | 0.608 | 0.633 |
| 190 | 1FSP_A | $\alpha/\beta$ | 124 | 0.843 | 0.847 | 0.816 | 0.858 | 0.843 | 0.846 |
| 191 | 1OOF_A | $\alpha/\beta$ | 124 | 0.731 | 0.770 | 0.628 | 0.628 | 0.756 | 0.794 |
| 192 | 1F98_A | $\alpha/\beta$ | 125 | 0.662 | 0.711 | 0.530 | 0.538 | 0.658 | 0.678 |
| 193 | 1FR0_A | $\alpha$ | 125 | 0.741 | 0.767 | 0.723 | 0.728 | 0.699 | 0.708 |
| 194 | 1HKQ_A | $\alpha/\beta$ | 125 | 0.623 | 0.688 | 0.645 | 0.645 | 0.610 | 0.633 |
| 195 | 2L74_A | $\alpha/\beta$ | 125 | 0.712 | 0.712 | 0.525 | 0.555 | 0.646 | 0.701 |
| 196 | 2P7L_A | $\alpha/\beta$ | 125 | 0.709 | 0.750 | 0.574 | 0.574 | 0.654 | 0.685 |
| 197 | 2RD5_D | $\alpha/\beta$ | 126 | 0.502 | 0.502 | 0.450 | 0.476 | 0.530 | 0.554 |
| 198 | 3BDB_A | $\alpha/\beta$ | 126 | 0.505 | 0.525 | 0.605 | 0.605 | 0.670 | 0.670 |
| 199 | 3CG4_A | $\alpha/\beta$ | 126 | 0.802 | 0.807 | 0.776 | 0.776 | 0.845 | 0.845 |
| 200 | 3CX5_G | $\alpha$ | 126 | 0.709 | 0.709 | 0.440 | 0.440 | 0.544 | 0.544 |
|  |  |  |  | First model | Best model | First model | Best model | First model | Best model |
| 201 | 5CJ3_B | $\alpha/\beta$ | 126 | 0.690 | 0.729 | 0.499 | 0.499 | 0.627 | 0.666 |
| 202 | 1B8Q_A | $\alpha/\beta$ | 127 | 0.436 | 0.437 | 0.244 | 0.268 | 0.506 | 0.521 |
| 203 | 1DBF_A | $\alpha/\beta$ | 127 | 0.661 | 0.661 | 0.553 | 0.553 | 0.639 | 0.648 |
| 204 | 1UFB_A | $\alpha$ | 127 | 0.788 | 0.788 | 0.751 | 0.751 | 0.801 | 0.801 |
| 205 | 2A9U_B | $\alpha$ | 127 | 0.783 | 0.794 | 0.664 | 0.664 | 0.719 | 0.738 |
| 206 | 3I9V_7 | $\alpha/\beta$ | 127 | 0.538 | 0.538 | 0.451 | 0.451 | 0.423 | 0.610 |
| 207 | 1DOI_A | $\alpha/\beta$ | 128 | 0.499 | 0.499 | 0.339 | 0.339 | 0.520 | 0.537 |
| 208 | 1GPQ_B | $\alpha/\beta$ | 128 | 0.640 | 0.641 | 0.470 | 0.470 | 0.664 | 0.680 |
| 209 | 1PXW_A | $\alpha/\beta$ | 128 | 0.655 | 0.720 | 0.536 | 0.536 | 0.757 | 0.776 |
| 210 | 4I60_A | $\alpha/\beta$ | 128 | 0.564 | 0.564 | 0.511 | 0.531 | 0.753 | 0.753 |
| 211 | 4JGX_B | $\alpha/\beta$ | 128 | 0.676 | 0.719 | 0.666 | 0.666 | 0.647 | 0.657 |
| 212 | 1AHK_A | $\beta$ | 129 | 0.483 | 0.483 | 0.552 | 0.552 | 0.512 | 0.581 |
| 213 | 2QVG_A | $\alpha/\beta$ | 129 | 0.794 | 0.821 | 0.795 | 0.795 | 0.810 | 0.810 |
| 214 | 2XGY_A | $\alpha$ | 129 | 0.624 | 0.632 | 0.543 | 0.543 | 0.436 | 0.589 |
| 215 | 1AX8_A | $\alpha/\beta$ | 130 | 0.576 | 0.589 | 0.456 | 0.471 | 0.428 | 0.563 |
| 216 | 1GQA_A | $\alpha/\beta$ | 130 | 0.763 | 0.791 | 0.738 | 0.738 | 0.756 | 0.796 |
| 217 | 2HYB_A | $\alpha/\beta$ | 130 | 0.618 | 0.625 | 0.616 | 0.616 | 0.772 | 0.772 |
| 218 | 4B0M_A | $\alpha/\beta$ | 131 | 0.554 | 0.605 | 0.434 | 0.495 | 0.510 | 0.510 |
| 219 | 1B4U_A | $\alpha$ | 132 | 0.616 | 0.619 | 0.642 | 0.642 | 0.592 | 0.611 |
| 220 | 3CAE_A | $\alpha/\beta$ | 132 | 0.508 | 0.508 | 0.328 | 0.328 | 0.471 | 0.471 |
| 221 | 4CXT_A | $\alpha/\beta$ | 132 | 0.703 | 0.703 | 0.457 | 0.516 | 0.706 | 0.706 |
| 222 | 1DCF_A | $\alpha/\beta$ | 133 | 0.822 | 0.823 | 0.763 | 0.763 | 0.823 | 0.823 |
| 223 | 1OA8_D | $\alpha/\beta$ | 133 | 0.295 | 0.320 | 0.241 | 0.261 | 0.276 | 0.309 |
| 224 | 1Y14_A | $\alpha$ | 133 | 0.442 | 0.571 | 0.410 | 0.445 | 0.508 | 0.554 |
| 225 | 4LE0_B | $\alpha/\beta$ | 133 | 0.895 | 0.907 | 0.800 | 0.870 | 0.914 | 0.914 |
| 226 | 4Z6J_A | $\beta$ | 133 | 0.488 | 0.507 | 0.276 | 0.486 | 0.643 | 0.643 |
| 227 | 1PMS_A | $\alpha/\beta$ | 135 | 0.533 | 0.568 | 0.544 | 0.544 | 0.614 | 0.614 |
| 228 | 1S56_B | $\alpha/\beta$ | 135 | 0.751 | 0.759 | 0.712 | 0.712 | 0.724 | 0.758 |
| 229 | 1OJG_A | $\alpha/\beta$ | 136 | 0.624 | 0.624 | 0.599 | 0.599 | 0.633 | 0.633 |
| 230 | 1SVJ_A | $\alpha/\beta$ | 136 | 0.750 | 0.756 | 0.639 | 0.639 | 0.757 | 0.765 |
| 231 | 1TWU_A | $\alpha/\beta$ | 137 | 0.738 | 0.738 | 0.585 | 0.585 | 0.606 | 0.606 |
| 232 | 1FJG_H | $\alpha/\beta$ | 138 | 0.712 | 0.740 | 0.470 | 0.470 | 0.782 | 0.782 |
| 233 | 3UE6_E | $\alpha/\beta$ | 138 | 0.685 | 0.690 | 0.572 | 0.582 | 0.797 | 0.797 |
| 234 | 1LNW_C | $\alpha/\beta$ | 139 | 0.812 | 0.826 | 0.769 | 0.769 | 0.663 | 0.752 |
| 235 | 4AIH_A | $\alpha/\beta$ | 139 | 0.772 | 0.772 | 0.652 | 0.666 | 0.643 | 0.682 |
| 236 | 1KQ6_A | $\alpha/\beta$ | 140 | 0.699 | 0.712 | 0.571 | 0.571 | 0.722 | 0.727 |
| 237 | 1NPB_A | $\alpha/\beta$ | 140 | 0.592 | 0.649 | 0.554 | 0.554 | 0.485 | 0.567 |
| 238 | 1OZ9_A | $\alpha/\beta$ | 141 | 0.677 | 0.700 | 0.600 | 0.621 | 0.807 | 0.807 |
| 239 | 1R5T_A | $\alpha/\beta$ | 141 | 0.691 | 0.711 | 0.625 | 0.625 | 0.739 | 0.739 |
| 240 | 2FA5_B | $\alpha/\beta$ | 142 | 0.714 | 0.730 | 0.658 | 0.687 | 0.625 | 0.648 |
| 241 | 2HQ7_B | $\alpha/\beta$ | 142 | 0.697 | 0.697 | 0.664 | 0.664 | 0.712 | 0.712 |
| 242 | 1HKX_E | $\alpha/\beta$ | 143 | 0.471 | 0.471 | 0.509 | 0.509 | 0.657 | 0.712 |
| 243 | 1HL6_D | $\alpha/\beta$ | 143 | 0.486 | 0.574 | 0.270 | 0.315 | 0.540 | 0.668 |
| 244 | 1S3J_A | $\alpha/\beta$ | 143 | 0.770 | 0.787 | 0.804 | 0.804 | 0.752 | 0.752 |
| 245 | 2F22_B | $\alpha/\beta$ | 143 | 0.671 | 0.685 | 0.566 | 0.566 | 0.737 | 0.771 |
| 246 | 4KA0_A | $\alpha/\beta$ | 143 | 0.798 | 0.804 | 0.708 | 0.708 | 0.849 | 0.849 |
| 247 | 5L8R_D | $\alpha/\beta$ | 143 | 0.435 | 0.464 | 0.287 | 0.318 | 0.360 | 0.428 |
| 248 | 1KSX_A | $\alpha/\beta$ | 144 | 0.652 | 0.697 | 0.537 | 0.537 | 0.730 | 0.730 |
| 249 | 1LE2_A | $\alpha$ | 144 | 0.267 | 0.267 | 0.299 | 0.380 | 0.357 | 0.457 |
| 250 | 1A3A_C | $\alpha/\beta$ | 146 | 0.607 | 0.643 | 0.493 | 0.555 | 0.758 | 0.758 |
|  |  |  |  | First model | Best model | First model | Best model | First model | Best model |
| 251 | 1K5D_B | $\alpha/\beta$ | 146 | 0.583 | 0.583 | 0.512 | 0.531 | 0.618 | 0.688 |
| 252 | 1LZW_B | $\alpha/\beta$ | 146 | 0.807 | 0.807 | 0.704 | 0.704 | 0.815 | 0.820 |
| 253 | 1HBG_A | $\alpha$ | 147 | 0.830 | 0.833 | 0.780 | 0.780 | 0.840 | 0.858 |
| 254 | 3E6M_E | $\alpha/\beta$ | 147 | 0.730 | 0.731 | 0.665 | 0.665 | 0.618 | 0.678 |
| 255 | 3PD2_A | $\alpha/\beta$ | 147 | 0.622 | 0.622 | 0.533 | 0.533 | 0.723 | 0.723 |
| 256 | 4GQY_A | $\alpha/\beta$ | 147 | 0.664 | 0.664 | 0.591 | 0.591 | 0.742 | 0.742 |
| 257 | 1UNG_D | $\alpha$ | 149 | 0.683 | 0.718 | 0.493 | 0.493 | 0.503 | 0.638 |
| 258 | 1GME_A | $\alpha/\beta$ | 150 | 0.528 | 0.528 | 0.250 | 0.375 | 0.478 | 0.485 |
| 259 | 1F1E_A | $\alpha/\beta$ | 151 | 0.667 | 0.667 | 0.374 | 0.432 | 0.476 | 0.551 |
| 260 | 2H30_A | $\alpha/\beta$ | 151 | 0.661 | 0.706 | 0.651 | 0.666 | 0.713 | 0.718 |
| 261 | 2PYB_A | $\alpha$ | 151 | 0.755 | 0.791 | 0.731 | 0.731 | 0.858 | 0.869 |
| 262 | 1NTV_A | $\alpha/\beta$ | 152 | 0.614 | 0.614 | 0.545 | 0.545 | 0.695 | 0.695 |
| 263 | 2NS9_B | $\alpha/\beta$ | 152 | 0.783 | 0.788 | 0.636 | 0.636 | 0.718 | 0.819 |
| 264 | 3GMX_A | $\alpha/\beta$ | 153 | 0.437 | 0.437 | 0.265 | 0.265 | 0.340 | 0.349 |
| 265 | 2GKC_A | $\alpha/\beta$ | 155 | 0.459 | 0.459 | 0.555 | 0.555 | 0.617 | 0.617 |
| 266 | 5JTM_A | $\alpha/\beta$ | 155 | 0.580 | 0.580 | 0.366 | 0.443 | 0.580 | 0.594 |
| 267 | 3ALU_A | $\alpha/\beta$ | 157 | 0.400 | 0.400 | 0.472 | 0.472 | 0.630 | 0.630 |
| 268 | 1OX7_A | $\alpha/\beta$ | 158 | 0.655 | 0.655 | 0.477 | 0.477 | 0.792 | 0.809 |
| 269 | 1USL_C | $\alpha/\beta$ | 158 | 0.756 | 0.775 | 0.671 | 0.671 | 0.775 | 0.790 |
| 270 | 2KBW_A | $\alpha$ | 160 | 0.722 | 0.756 | 0.583 | 0.659 | 0.693 | 0.725 |
| 271 | 3F8L_A | $\alpha/\beta$ | 162 | 0.628 | 0.654 | 0.504 | 0.541 | 0.863 | 0.868 |
| 272 | 1C03_A | $\alpha/\beta$ | 163 | 0.645 | 0.645 | 0.515 | 0.542 | 0.679 | 0.700 |
| 273 | 3H05_B | $\alpha/\beta$ | 163 | 0.605 | 0.605 | 0.442 | 0.442 | 0.723 | 0.764 |
| 274 | 4NBI_A | $\alpha/\beta$ | 163 | 0.503 | 0.503 | 0.524 | 0.524 | 0.683 | 0.704 |
| 275 | 1FW9_A | $\alpha/\beta$ | 164 | 0.552 | 0.552 | 0.432 | 0.447 | 0.607 | 0.634 |
| 276 | 2AEN_A | $\alpha/\beta$ | 164 | 0.421 | 0.421 | 0.313 | 0.313 | 0.261 | 0.396 |
| 277 | 1C41_A | $\alpha/\beta$ | 165 | 0.696 | 0.696 | 0.706 | 0.706 | 0.712 | 0.712 |
| 278 | 1F3Y_A | $\alpha/\beta$ | 165 | 0.340 | 0.368 | 0.433 | 0.433 | 0.513 | 0.548 |
| 279 | 1S2D_A | $\alpha/\beta$ | 165 | 0.642 | 0.695 | 0.602 | 0.605 | 0.721 | 0.721 |
| 280 | 2LRB_A | $\alpha/\beta$ | 165 | 0.607 | 0.607 | 0.585 | 0.585 | 0.679 | 0.679 |
| 281 | 3V1O_A | $\alpha/\beta$ | 165 | 0.595 | 0.595 | 0.510 | 0.510 | 0.662 | 0.662 |
| 282 | 1PGV_A | $\alpha/\beta$ | 167 | 0.782 | 0.806 | 0.584 | 0.592 | 0.788 | 0.788 |
| 283 | 2FKB_C | $\alpha/\beta$ | 167 | 0.522 | 0.522 | 0.482 | 0.493 | 0.635 | 0.635 |
| 284 | 1AP7_A | $\alpha$ | 168 | 0.775 | 0.787 | 0.590 | 0.590 | 0.736 | 0.736 |
| 285 | 2C4W_A | $\alpha/\beta$ | 168 | 0.636 | 0.649 | 0.661 | 0.661 | 0.723 | 0.723 |
| 286 | 2O70_F | $\alpha$ | 168 | 0.750 | 0.774 | 0.607 | 0.607 | 0.782 | 0.782 |
| 287 | 2WGP_A | $\alpha/\beta$ | 168 | 0.762 | 0.769 | 0.662 | 0.668 | 0.835 | 0.844 |
| 288 | 3M1N_B | $\alpha/\beta$ | 168 | 0.433 | 0.439 | 0.371 | 0.385 | 0.492 | 0.501 |
| 289 | 1XJA_C | $\alpha/\beta$ | 169 | 0.643 | 0.654 | 0.549 | 0.570 | 0.691 | 0.697 |
| 290 | 1YG2_A | $\alpha/\beta$ | 169 | 0.445 | 0.623 | 0.393 | 0.448 | 0.438 | 0.451 |
| 291 | 1QZG_A | $\alpha/\beta$ | 170 | 0.561 | 0.561 | 0.387 | 0.387 | 0.687 | 0.687 |
| 292 | 1NF6_F | $\alpha/\beta$ | 171 | 0.806 | 0.806 | 0.703 | 0.718 | 0.815 | 0.842 |
| 293 | 1NQZ_A | $\alpha/\beta$ | 171 | 0.605 | 0.607 | 0.452 | 0.523 | 0.677 | 0.677 |
| 294 | 5IAO_A | $\alpha/\beta$ | 171 | 0.491 | 0.493 | 0.470 | 0.548 | 0.690 | 0.696 |
| 295 | 6AQ3_B | $\alpha/\beta$ | 171 | 0.558 | 0.569 | 0.489 | 0.493 | 0.508 | 0.593 |
| 296 | 1KOH_D | $\alpha/\beta$ | 172 | 0.752 | 0.752 | 0.597 | 0.597 | 0.544 | 0.553 |
| 297 | 1TEO_A | $\alpha/\beta$ | 173 | 0.554 | 0.554 | 0.545 | 0.545 | 0.647 | 0.647 |
| 298 | 2A5Y_A | $\alpha$ | 173 | 0.698 | 0.723 | 0.550 | 0.550 | 0.623 | 0.719 |
| 299 | 1AK6_A | $\alpha/\beta$ | 174 | 0.514 | 0.514 | 0.507 | 0.507 | 0.630 | 0.630 |
| 300 | 1QFT_A | $\alpha/\beta$ | 175 | 0.506 | 0.506 | 0.534 | 0.562 | 0.689 | 0.689 |
|  |  |  |  | First model | Best model | First model | Best model | First model | Best model |
| 301 | 2L5P_A | $\alpha/\beta$ | 175 | 0.632 | 0.632 | 0.595 | 0.595 | 0.658 | 0.676 |
| 302 | 2C2F_A | $\alpha/\beta$ | 178 | 0.802 | 0.802 | 0.747 | 0.758 | 0.802 | 0.802 |
| 303 | 1NGL_A | $\alpha/\beta$ | 179 | 0.552 | 0.552 | 0.526 | 0.557 | 0.602 | 0.602 |
| 304 | 3IAM_2 | $\alpha/\beta$ | 179 | 0.530 | 0.616 | 0.345 | 0.382 | 0.509 | 0.511 |
| 305 | 2Z3B_A | $\alpha/\beta$ | 180 | 0.751 | 0.751 | 0.443 | 0.443 | 0.661 | 0.661 |
| 306 | 1Y1X_A | $\alpha/\beta$ | 182 | 0.667 | 0.667 | 0.738 | 0.738 | 0.629 | 0.648 |
| 307 | 1RZ3_A | $\alpha/\beta$ | 184 | 0.463 | 0.515 | 0.516 | 0.597 | 0.604 | 0.604 |
| 308 | 1WLQ_C | $\alpha/\beta$ | 185 | 0.629 | 0.648 | 0.501 | 0.501 | 0.576 | 0.593 |
| 309 | 1TJF_B | $\alpha/\beta$ | 186 | 0.791 | 0.791 | 0.726 | 0.746 | 0.772 | 0.782 |
| 310 | 3O61_A | $\alpha/\beta$ | 187 | 0.348 | 0.418 | 0.407 | 0.434 | 0.567 | 0.567 |
| 311 | 1TLJ_A | $\alpha/\beta$ | 189 | 0.536 | 0.536 | 0.527 | 0.527 | 0.470 | 0.554 |
| 312 | 1DTP_A | $\alpha/\beta$ | 190 | 0.222 | 0.258 | 0.160 | 0.188 | 0.223 | 0.314 |
| 313 | 1F15_C | $\alpha/\beta$ | 191 | 0.197 | 0.222 | 0.149 | 0.162 | 0.215 | 0.273 |
| 314 | 1GXD_C | $\alpha/\beta$ | 192 | 0.455 | 0.455 | 0.197 | 0.255 | 0.427 | 0.484 |
| 315 | 1VCY_A | $\alpha/\beta$ | 193 | 0.464 | 0.464 | 0.351 | 0.355 | 0.322 | 0.411 |
| 316 | 2VUL_A | $\alpha/\beta$ | 193 | 0.411 | 0.411 | 0.393 | 0.398 | 0.424 | 0.645 |
| 317 | 4ZBY_A | $\alpha/\beta$ | 194 | 0.657 | 0.657 | 0.591 | 0.591 | 0.794 | 0.794 |
| 318 | 1IOO_A | $\alpha/\beta$ | 196 | 0.447 | 0.447 | 0.404 | 0.404 | 0.530 | 0.574 |
| 319 | 5EKT_A | $\alpha/\beta$ | 196 | 0.461 | 0.461 | 0.525 | 0.528 | 0.690 | 0.713 |
| 320 | 4K1F_A | $\alpha/\beta$ | 198 | 0.535 | 0.601 | 0.529 | 0.572 | 0.785 | 0.785 |

**Table S2:** Prediction results (TM-score) of MMpred, C-QUARK, RaptorX-DeepModeller, BAKER- ROSETTASERVER and MULTICOM_ CLUSTER for 24 free modeling (FM) targets of CASP13.

| No. | PDB | Size | MMpred | C-QUARK | RaptorX-Deep<br>Modeller | BAKER-ROSETTAS<br>ERVER | MULTICOM<br>_ CLUSTER |
| --- | --- | --- | --- | --- | --- | --- | --- |
| 1 | T0955-D1 | 41 | 0.67 | 0.73 | 0.59 | \ | 0.77 |
| 2 | T0953s2-D1 | 44 | 0.32 | 0.36 | 0.18 | 0.33 | 0.20 |
| 3 | T0953s1-D1 | 67 | 0.38 | 0.40 | 0.28 | 0.19 | 0.38 |
| 4 | T0990-D1 | 76 | 0.44 | 0.58 | 0.40 | 0.37 | 0.36 |
| 5 | T0958-D1 | 77 | 0.73 | 0.55 | 0.66 | 0.53 | 0.55 |
| 6 | T1008-D1 | 77 | 0.70 | 0.35 | 0.28 | 0.56 | 0.38 |
| 7 | T0963-D2 | 82 | 0.43 | 0.46 | 0.48 | 0.36 | 0.28 |
| 8 | T0960-D2 | 84 | 0.42 | 0.44 | 0.49 | 0.28 | 0.38 |
| 9 | T0970-D1 | 85 | 0.54 | 0.51 | 0.54 | 0.40 | 0.33 |
| 10 | T0953s2-D3 | 93 | 0.16 | 0.35 | 0.29 | 0.19 | 0.14 |
| 11 | T1021s3-D2 | 97 | 0.42 | 0.45 | 0.59 | 0.19 | 0.27 |
| 12 | T0980s1-D1 | 104 | 0.45 | 0.54 | 0.37 | 0.41 | 0.25 |
| 13 | T0957s1-D1 | 108 | 0.42 | 0.40 | 0.37 | 0.42 | 0.31 |
| 14 | T0953s2-D2 | 111 | 0.50 | 0.48 | 0.69 | 0.47 | 0.22 |
| 15 | T0968s2-D1 | 115 | 0.60 | 0.65 | 0.59 | 0.66 | 0.42 |
| 16 | T0968s1-D1 | 118 | 0.69 | 0.56 | 0.62 | 0.74 | 0.43 |
| 17 | T0957s2-D1 | 155 | 0.70 | 0.53 | 0.65 | 0.48 | 0.51 |
| 18 | T1022s1-D1 | 156 | 0.37 | 0.55 | 0.58 | 0.40 | 0.38 |
| 19 | T1021s3-D1 | 166 | 0.43 | 0.64 | 0.66 | 0.50 | 0.50 |
| 20 | T0990-D3 | 213 | 0.25 | 0.22 | 0.22 | 0.23 | 0.24 |
| 21 | T0990-D2 | 231 | 0.40 | 0.37 | 0.35 | 0.26 | 0.25 |
| 22 | T1005-D1 | 326 | 0.43 | 0.70 | 0.70 | 0.70 | 0.69 |
| 23 | T0950-D1 | 342 | 0.41 | 0.44 | 0.56 | 0.46 | 0.22 |
| 24 | T0969-D1 | 354 | 0.33 | 0.64 | 0.65 | 0.49 | 0.44 |

**Table S3:**
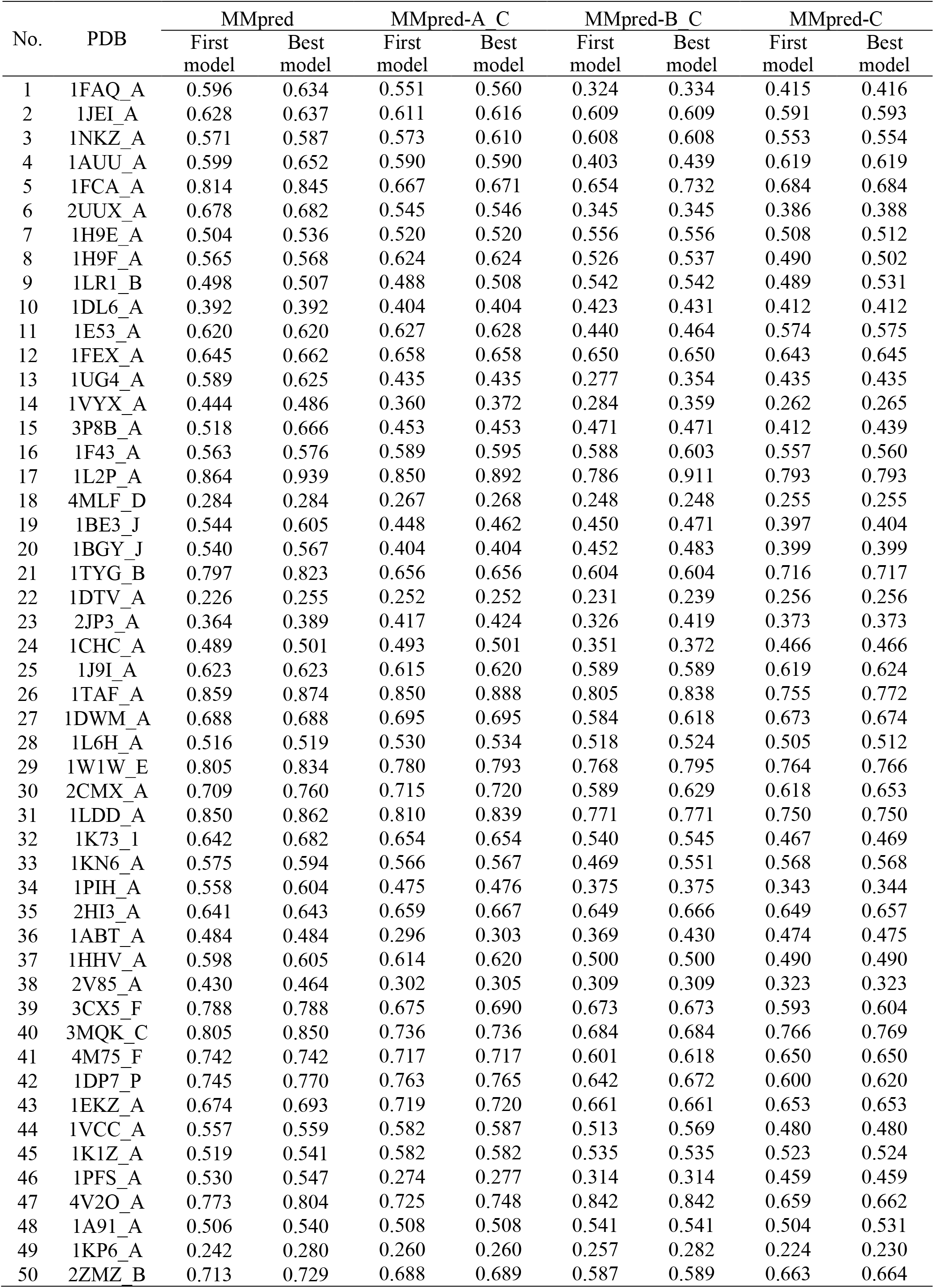
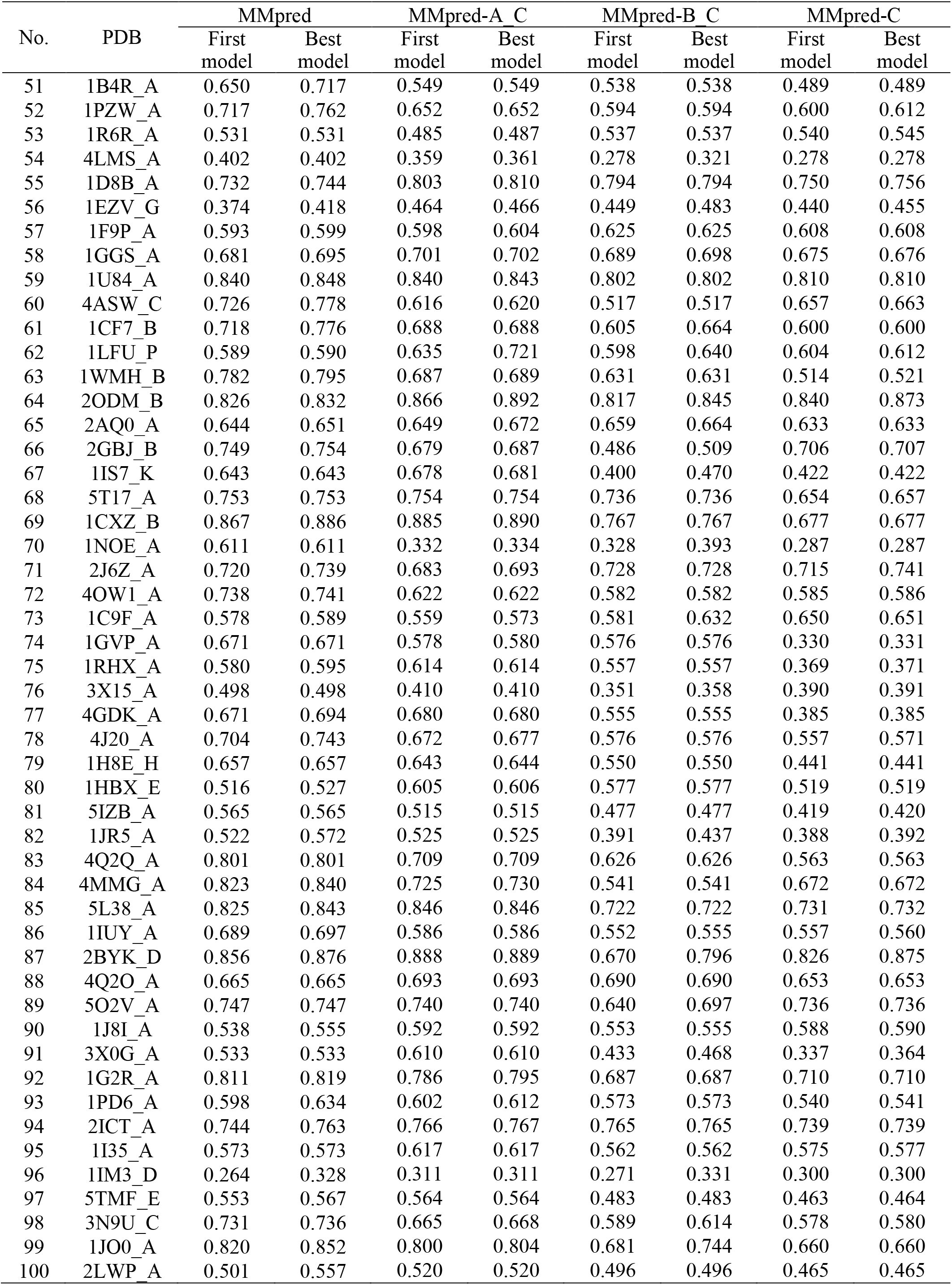
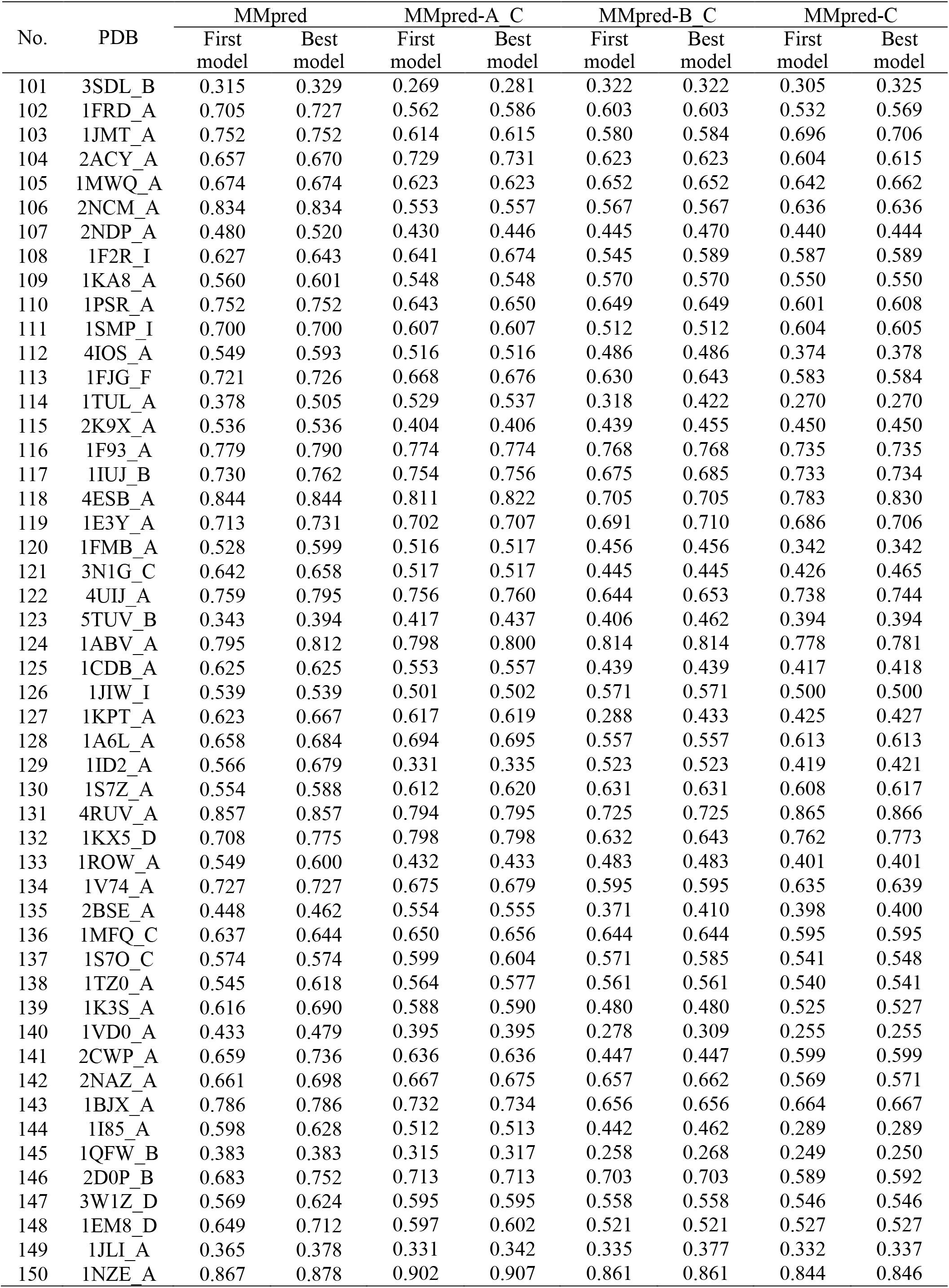
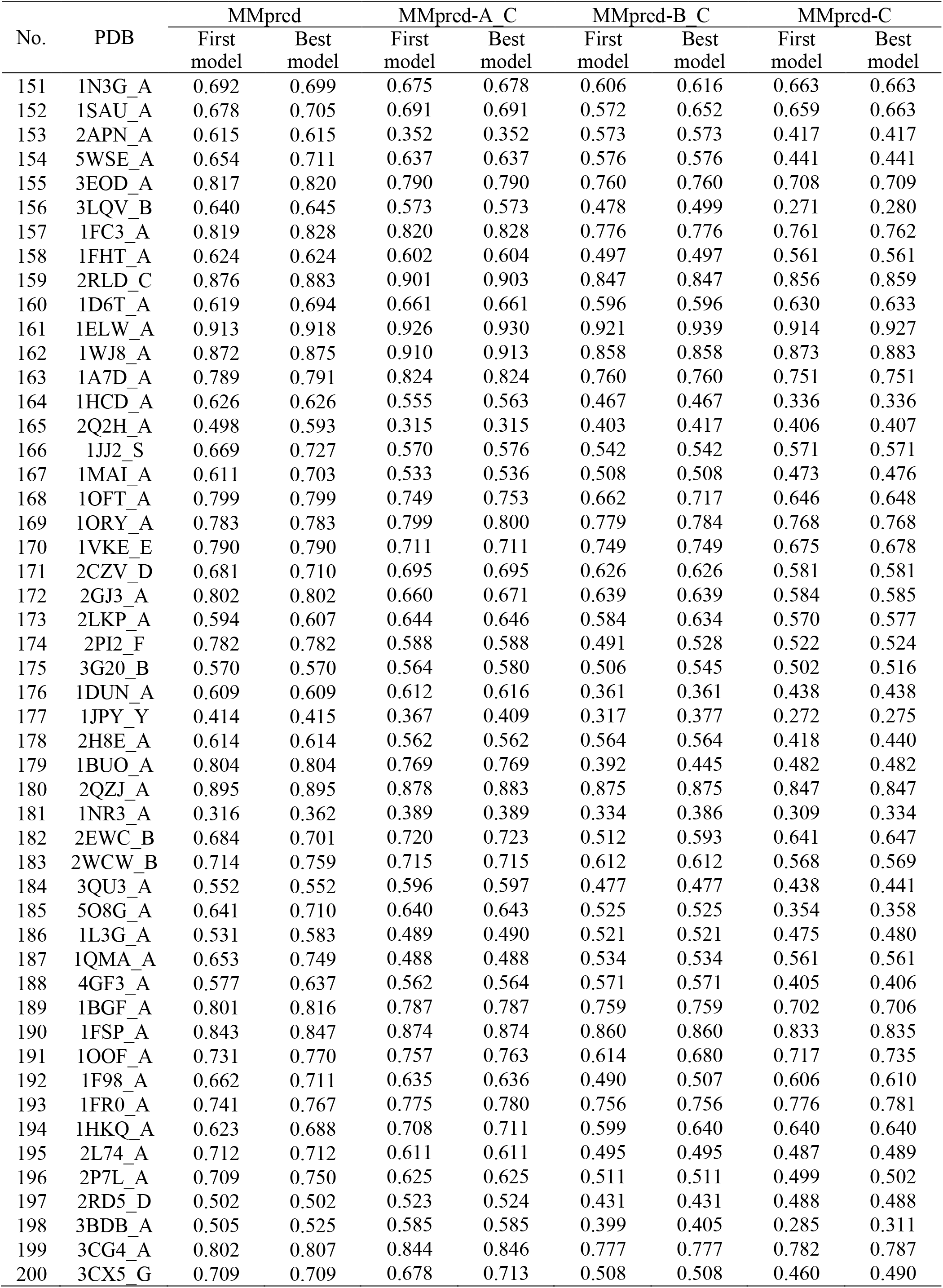
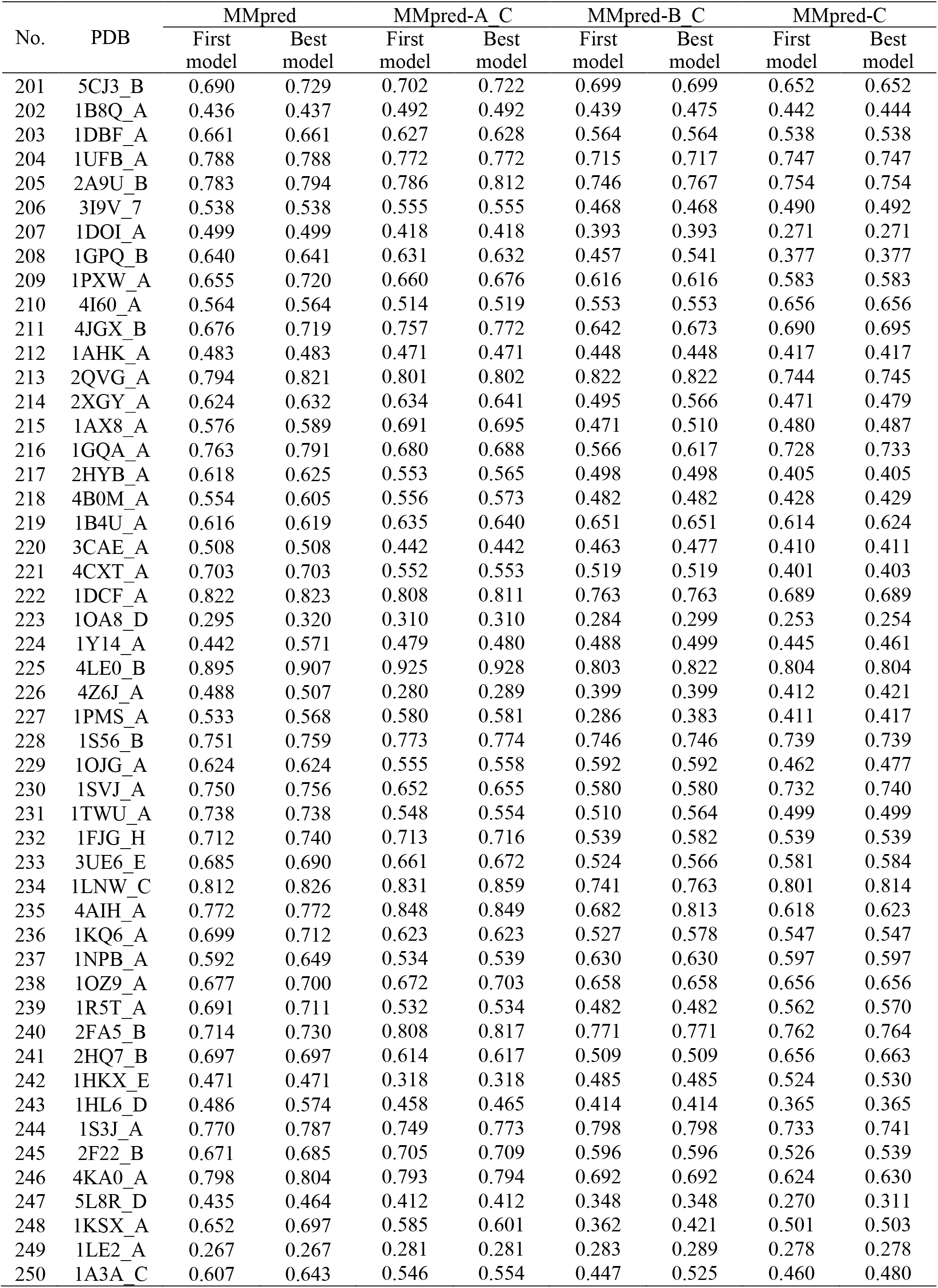
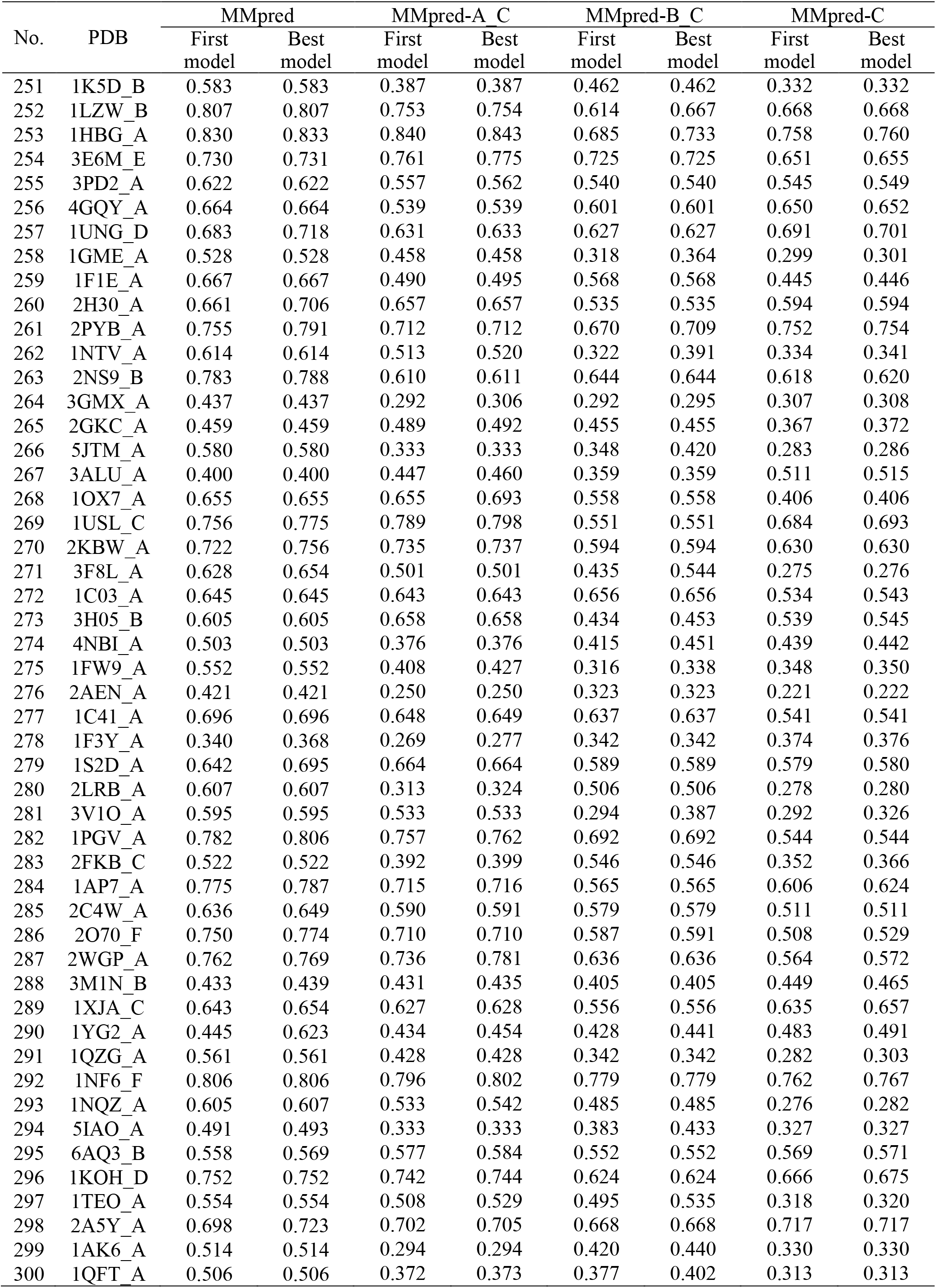
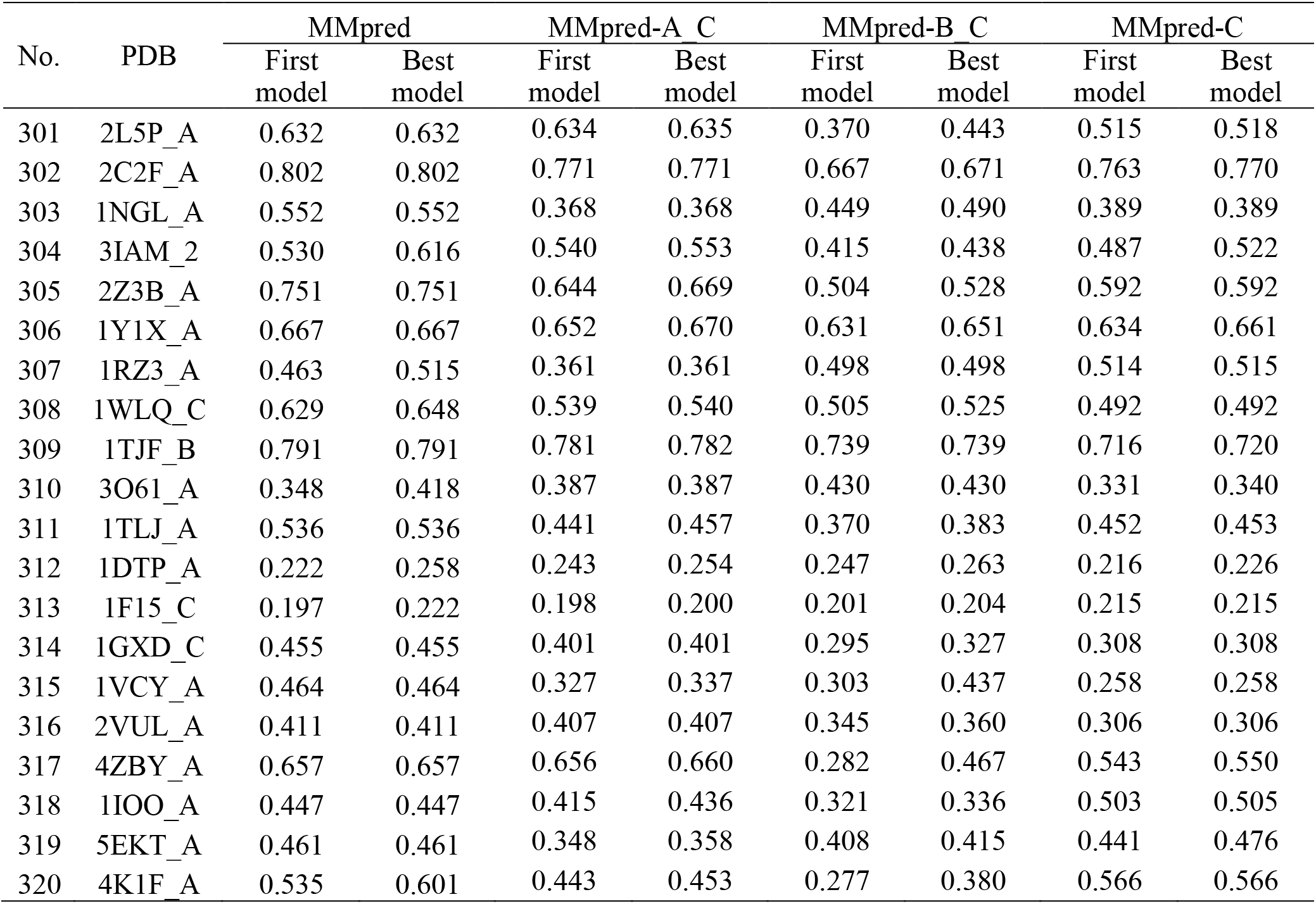
Prediction results (TM-score) of MMpred, MMpred-A_C, MMpred-B_C and MMpred-C for 320 benchmark proteins.

## References

Abriata, L.A., Tamò, G.E. and Dal Peraro, M. (2019) A further leap of improvement in tertiary structure prediction in CASP13 prompts new routes for future assessments. Proteins, 87, 1100–1112.

Bradley, P., Misura, K.M.S. and Baker, D. (2005) Toward high resolution de novo structure prediction for small proteins. Science, 309, 1868–1871.

Brunger, A.T. (2007) Version 1.2 of the crystallography and NMR system. Nat. Protoc., 2, 2728–2733.

Bowman, G.R. and Pande, V.S. (2009) Simulated tempering yields insight into the low-resolution Rosetta scoring functions. Proteins, 74, 777–788.

Cheng, J.L., Choe, M.H., Elofsson, A., et al. (2019) Estimation of model accuracy in CASP13. Proteins, 87, 1361–1377.

De Jong, K.A. (1975) An analysis of the behaviore of a class of genetic adaptive systems. Doctoral Dissertation, University of Michigan.

Dill, K.A. and MacCallum, J.L. (2012) The protein-folding problem, 50 years on. science, 338, 1042–1046.

Francois, B., Zhou, Y., Shrestha, R., and Zhang, K.Y.J. (2011) Entropy-accelerated exact clustering of protein decoys. Bioinformatics, 27, 939–945.

Fu, L., Niu, B.F., Zhu, Z.W., Wu, S.T. and Li, W.Z. (2012) CD-HIT: accelerated for clustering the next-generation sequencing data. Bioinformatics, 28, 3150–3152.

Fox, N.K., Brenner, S.E., and Chandonia, J.M. (2014) SCOPe: structural Classification of Proteins—extended, integrating SCOP and ASTRAL data and classification of new structures. Nucleic Acids Res, 42, D304–D309.

Greener, J.G., Kandathil, S.M. and Jones, D.T. (2019) Deep learning extends de novo protein modelling coverage of genomes using iteratively predicted structural constraints. Nature Communications, 10, 3977.

Garza-Fabre, M., Kandathil, S.M., Handl, J., et al. (2016) Generating, Maintaining, and Exploiting Diversity in a Memetic Algorithm for Protein Structure Prediction. Evolutionary Computation, 24, 577–607.

Hou, J., Wu, T.Q., Cao, R.Z. and Cheng, J.L. (2019) Protein tertiary structure modeling driven by deep learning and contact distance prediction in CASP13. Proteins, 87, 1165–1178.

Hubbard, T.J.P. (1999) RMS/coverage graphs: a qualitative method for comparing three-dimensional protein structure predictions, Proteins, Suppl 3(S3), 15–21.

Kandathil, S.M., Greener, J.G. and Jones, D.T. (2019) Prediction of interresidue contacts with DeepMetaPSICOV in CASP13. Proteins, 87, 1092–1099.

Kuhlman, B. and Bradley, P. (2019) Advances in protein structure prediction and design. Nature Reviews Molecular Cell Biology, 20.

Kaufman, L. and Rousseeuw, P.J. (2009) Finding Groups in Data: An Introduction to Cluster Analysis. Wiley-Interscience, New York.

Lazaridis, T. and Karplus, M. (2000) Effective energy functions for protein structure prediction. Current Opinion in Structural Biology, 10, 139–145.

Liu, J., Zhou, X.G., Zhang, Y. and Zhang, G.J. (2019) CGLFold: a contact-assisted de novo protein structure prediction using global exploration and loop perturbation sampling algorithm. Bioinformatics, 36, 2443–2450.

Lindorff-Larsen, K., Piana, S., Dror, R.O., and Shaw, D.E. (2011) How fast-folding proteins fold. Science, 334, 517–520.

Ling, Q., Wu, G., Yang, Z.Y. and Wang, Q.P. (2008) Crowding clustering genetic algorithm for multimodal function optimization. Applied Soft Computing, 8, 88–95.

Li, Z. and Scheraga, H.A. (1987) Monte Carlo-minimization approach to the multiple minima problem in protein folding. Proceedings of the National Academy of Sciences of the United States of America, 84, 6611–6615.

Li, Y., Hu, J., Zhang, C.X., Yu, D.J. and Zhang, Y. (2019) ResPRE: high-accuracy protein contact prediction by coupling precision matrix with deep residual neural networks. Bioinformatics, 35, 4647–4655.

Mariani, V., Biasini, M., Barbato, A. and Schwede, T. (2013) lDDT: a local superposition-free score for comparing protein Structures and models using distance difference tests. Bioinformatics, 29, 2722–2728.

Olson, B. and Shehu, A. (2014) Multi-objective optimization techniques for conformational sampling in template-free protein structure prediction. International Conference on Bioinf and Comp Biol (BICoB), 2.

Ovchinnikov, S., Park, H., Kim, D.E., DiMaio, F. and Baker, D. (2018) Protein structure prediction using rosetta in casp12. Proteins, 86, 113–121.

Park, H., Ovchinnikov, S., Kim, D.E., DiMaio, F. and Baker, D. (2018) Protein homology model refinement by large-scale energy optimization. Proceedings of the National Academy of Sciences of the United States of America, 115, 3054–3059.

Park, H., Lee, G.R., Kim, D.E., Anishchenko, I., Cong, Q. and Baker, D. (2019) High-accuracy refinement using Rosetta in CASP13. Proteins, 87, 1276–1282.

Peng, C.X. Zhou, X.G. and Zhang, G.J. (2020) De novo Protein Structure Prediction by Coupling Contact with Distance Profile. IEEE/ACM Transactions on Computational Biology and Bioinformatics, doi:10.1109/TCBB.2020.3000758.

Rohl, C.A., Strauss, C.E.M., Misura, K.M.S. and Baker, D. (2004) Protein structure prediction using Rosetta. Methods in Enzymology. Methods in Enzymology, 383, 66–93.

Rodriguez, A. and Laio, A. (2014) Clustering by fast search and find of density peaks. Science, 334, 1492–1496.

Raman, S., Qian, B., Baker, D. and Walker, R.C. (2008) Advances in Rosetta protein structure prediction on massively parallel systems. IBM Journal of Research and Development, 52, 7–17.

Senior, A.W., Evans, R., Jumper, J., Kirkpatrick, J., Sifre, L. Green, T., et al. (2020) Improved protein structure prediction using potentials from deep learning. Nature, 577, 706–710.

Senior, A.W., Evans, R., Jumper, J., Kirkpatrick, J., Sifre, L., Green, T., et al. (2019) Protein structure prediction using multiple deep neural networks in the 13^th^ Critical Assessment of Protein Structure Prediction (CASP13). Proteins, 87, 1141–1148.

Stoean, C., Preuss, M., Stoean, R. and Dumitrescu, D. (2010) Multimodal optimization by means of a topological species conservation algorithm. IEEE Transactions on Evolutionary Computation, 14, 842–864.

Shrestha, R., Fajardo, E., Gil, N., Fidelis, K., Kryshtafovych, A., Monastyrskyy, B. and Fiser, A. (2019) Assessing the accuracy of contact predictions in CASP13. Proteins, 87, 1058–1068.

Simoncini, D., Schiex, T. and Zhang, K.Y.J. (2017) Balancing exploration and exploitation in population-based sampling improves fragment-based de novo protein structure prediction. Proteins, 85, 852–858.

Sareni, B. and Krahenbuhl, L. (1998) Fitness sharing and niching methods revisited. IEEE Transactions on Evolutionary Computation, 2, 97–106.

Thomsen, R. (2004) Multimodal optimization using crowding-based differential evolution. In: IEEE Congress on Evolutionary Computation, 1382–1389.

Wang, S., Sun, S.Q., Li, Z., Zhang, R.Y., and Xu, J.B. (2017) Accurate De novo Prediction of Protein Contact Map by Ultra-Deep Learning Model. Plos Computational Biology, 13, e1005324.

Xu, J.B. (2019) Distance-based protein folding powered by deep learning. Proceedings of the National Academy of Sciences of the United States of America, 116, 16856–16865.

Xu, J.B. and Wang, S. (2019) Analysis of distance-based protein structure prediction by deep learning in CASP13. Proteins, 87, 1069–1081.

Xu, D. and Zhang, Y. (2012) Ab initio protein structure assembly using continuous structure fragments and optimized knowledge-based force field. Proteins, 80, 1715–1735.

Yang, J.Y., Anishchenko, I., Park, H., Peng, Z.L, Ovchinnikov, S. and Baker, D. (2020) Improved protein structure prediction using predicted interresidue orientations. Proceedings of the National Academy of Sciences of the United States of America, 117, 1496–1503.

Zheng, W., Li, Y., Zhang, C.X., Pearce, R., Mortuza, S.M. and Zhang, Y. (2019) Deep-learning contact-map guided protein structure prediction in CASP13. Proteins, 87, 1149–1164.

Zhang, Y., Kihara, D. and Skolnick, J. (2002) Local energy landscape flattening: parallel hyperbolic Monte Carlo sampling of protein folding. Proteins, 48, 192–201.

Zhang, Y. and Skolnick, J. (2004) Scoring function for automated assessment of protein structure template quality. Proteins, 57, 702–710.

Zhang, Y. and Skolnick, J. (2004) SPICKER: a clustering approach to identify near-native protein folds. Journal of Computational Chemistry, 25, 865–871.

Zhou, X.G., Zhang, G.J., Hao, X.H., Xu, D.W. and Yu, L. (2016) Enhanced differential evolution using local Lipschitz underestimate strategy for computationally expensive optimization problems. Applied Soft Computing, 48, 169–181.

Zhou, X.G. and Zhang, G.J. (2017) Abstract convex underestimation assisted multistage differential evolution. IEEE transactions on cybernetics, 47, 2730–2741.

Zhou, X.G., Hu, J., Zhang. C.X., Zhang, G.J. and Zhang, Y. (2019) Assembling multidomain protein structures through analogous global structural alignments. Proceedings of the National Academy of Sciences of the United States of America, 116, 15930–15938.

Zhou, X.G. and Zhang, G.J. (2019) Differential evolution with underestimation-based multimutation strategy. IEEE Transactions on Cybernetics, 49, 1353–1364.

Zhou, X.G., Peng, C.X., Liu, J., Zhang, Y., and Zhang, G.J. (2020) Underestimation-assisted global-local cooperative differential evolution and the application to protein structure prediction. IEEE Transactions on Evolutionary Computation., 24, 536–550.

